# Empirical Bayes Estimation of Sample-Specific Correlation Networks for Cross-Sectional and Longitudinal Omics Data

**DOI:** 10.64898/2026.10.08.757613

**Authors:** Enakshi Saha, Hui Chen, Rahul Ghosal

## Abstract

Sample-specific molecular networks, which capture how an individual’s gene regulatory structure may deviate from the population as a whole, are important tools for understanding heterogeneity in health and disease. However, most existing methods either do not accommodate uncertainty quantification or cannot handle missing data, non-Gaussian modalities, and longitudinal structure common in modern multi-omic studies. For example, copy number alterations in cancer genomic studies are non-Gaussian and often only partially observed, and their regulatory effects on gene expression may vary substantially across patients, variation that a single, population-averaged network cannot capture. To address these challenges, we propose PRISM (Personalized Relational Inference via Sample-specific Modeling), an empirical Bayes framework for estimating sample-specific covariance matrices from cross-sectional and longitudinal multi-omic data. PRISM models each individual sample’s covariance matrix as drawn from an inverse-Wishart prior whose parameters are estimated from the data and allowed to vary across subpopulations or evolve over time, yielding a closed-form posterior and edge-level uncertainty quantification, ensuring scalability to high-dimensional omics data. In simulation experiments, PRISM achieves point-estimate accuracy comparable to or better than existing methods, substantially improved uncertainty calibration under non-Gaussian marginals, and lower computational cost, while maintaining robust coverage under moderate missingness. We apply PRISM to a yeast cell-cycle time course dataset, recovering time-varying biologically meaningful regulatory relationships independently documented in literature, and to paired copy number and gene expression data from ovarian cancer patients, identifying copy-number-expression interactions associated with survival, including TFs HNF4G and PAX8 which have independent evidence of clinical relevance in this cancer.

## 1 Introduction

Health and disease are rarely driven by a single gene in isolation but rather by a network of interacting genes and molecular regulators (Barabási et al. 2011). For example, high-grade serous ovarian carcinoma is heavily influenced by the interactions between TF copy number alterations (CNA) and the genes targeted by these TFs. It is among the most copy-number-driven cancers profiled by The Cancer Genome Atlas (TCGA), and patients with the same clinical diagnosis and stage often experience drastically different disease trajectories and therapy response (Network et al. 2011). A central hypothesis in precision oncology is that this variability is driven by differences in how TFs regulate their target genes from one patient’s tumor to another (Collins & Varmus 2015), so that a single, population-level regulatory network computed by most existing multi-omic network inference methods (Zhang et al. 2005, Danaher et al. 2014, Ha et al. 2021, Chakraborty et al. 2024) obscures the heterogeneity underlying individual risk, progression, and therapy response. Estimating a separate, patient-specific network relating each TF’s copy-number dosage to its target genes’ expression, and distinguishing which of these patient-specific regulatory associations are genuine from those that arise by chance, is essential for selecting a precision treatment based on how a tumor’s regulatory circuitry behaves in that patient, rather than on a population-average relationship that may not hold for them. Sample-specific networks of this kind have proven useful for identifying patient-specific regulatory drivers of disease (Saha et al. 2024, Chen, Saha, Fischer, Ben Guebila, Fanfani, Shutta, Padi, Glass, DeMeo, Lopes-Ramos et al. 2025, Saha et al. 2025), characterizing cell-to-cell heterogeneity in single-cell data (Dai et al. 2019, Wang et al. 2021), and tracking how regulatory structure evolves over developmental or disease time courses (Bodein et al. 2022).

This raises three challenges common to modern multi-omic cancer cohorts. First, CNA data are non-Gaussian, with a point mass at the diploid copy number alongside a continuous tail for gains and losses (Figure 1), so methods assuming Gaussian marginals risk distorted estimates. Second, patients are often missing a modality, or have partial missingness within a modality. Third, distinguishing a genuine regulatory association from one arising by chance requires a calibrated per-edge uncertainty measure to recover sparse, biologically plausible networks. Moreover, these challenges are not confined to a single snapshot in time: existing sample-specific network methods are designed for cross-sectional data and were never built for samples collected longitudinally, for instance across phases of a cell cycle or stages of disease progression, where an individual’s own regulatory structure may itself change.

**Figure 1:**
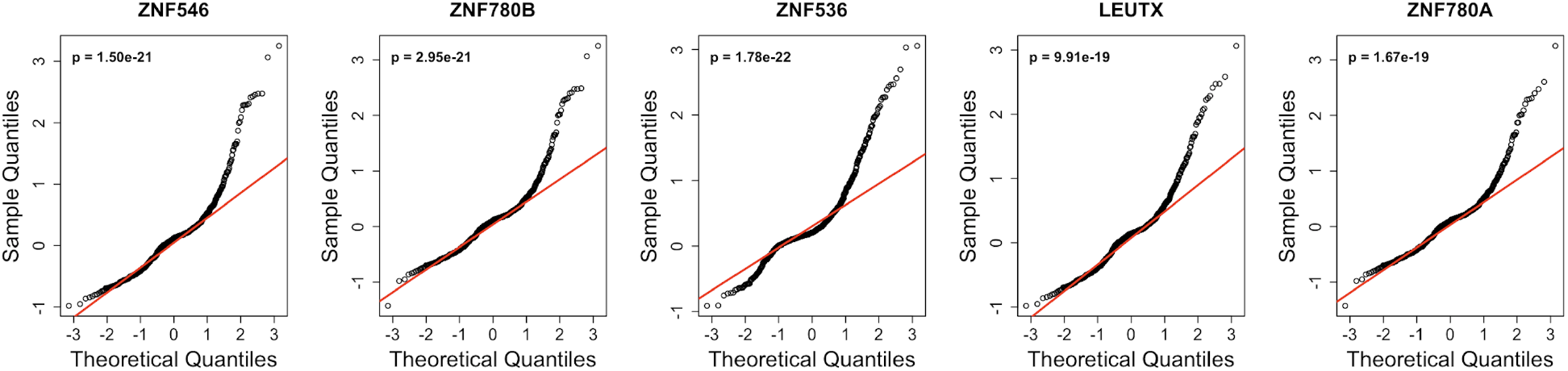
QQ-plots against a standard normal distribution for the five TFs with the highest CNA variance in TCGA Ovarian Cancer data, with the Shapiro–Wilk test *p*-value reported in each panel. All five show a clear departure from normality.

Several methods have been proposed to estimate sample-specific molecular networks, but none resolves these problems together. Perturbation- or interpolation-based methods (Kuijjer et al. 2019, Liu et al. 2016, Dai et al. 2019) give a point estimate per individual but no edge-level uncertainty. SWEET (Chen et al. 2023) tests each edge against the pooled distribution of all edges and individuals, a generic outlier test that ignores each edge’s baseline variability. Recent empirical Bayes models tackle this with a hierarchical shrinkage framework: BONOBO (Saha et al. 2024) estimates sample-specific covariance (coexpression) matrices, capturing marginal dependence between features, while SIREN (Saha 2026) estimates sample-specific partial correlations under a Gaussian graphical model, capturing conditional dependence instead. Both offer principled uncertainty quantification, but both are limited to Gaussian data without missingness and were built only for cross-sectional data.

We propose PRISM (*Personalized Relational Inference via Sample-specific Modeling*), a scalable empirical Bayes framework that addresses the challenges mentioned above and in addition, extends to discrete population heterogeneity, where the population consists of several subpopulations whose members vary only mildly from each other but substantially from members of other subpopulations, such as populations of different ancestry. In every extension PRISM estimates a covariance matrix for every sample in either cross-sectional or longitudinal data, where the sample-specific covariance matrices are assumed to follow a conjugate inverse-Wishart prior distribution whose parameters are estimated from the data, thus borrowing strength across samples and yielding a closed-form posterior. For longitudinal data, the prior itself evolves over time, so that each sample’s prior is informed only by nearby time points rather than the full cohort.

### Summary of our contributions

Our approach makes the following contributions:

#### Methodological contributions

PRISM provides a computationally efficient empirical Bayes framework for sample-specific covariance network estimation and further extends this to handle missing data, non-Gaussian modalities through a Gaussian copula, discrete population heterogeneity through group-specific priors, and longitudinal data through a prior that evolves over time, while retaining a closed-form posterior for every extension. PRISM also provides edge-level credible intervals, both exact and via a fast closed-form approximation, with a calibration that prevents the anti-conservative intervals that arise in empirical Bayes inference (Morris 1983, Laird & Louis 1987). Through simulation experiments, we show that PRISM gives comparable or more accurate point estimates and superior interval estimates with better coverage than existing methods for both cross-sectional and longitudinal data, remains robust under missing data, and is computationally more efficient than existing cross-sectional counterparts.

#### Scientific contributions

We apply PRISM to a yeast cell-cycle longitudinal dataset and recover cell-cycle regulatory relationships independently documented in the literature, reproducibly across biological replicates. We then apply PRISM to paired copy number and expression data from TCGA ovarian tumors to search for copy-number-driven regulatory interactions predictive of survival. Among the edges we identify, *HNF4G*-*GTF2H5* and *PAX8* -*CDHR5* emerge as candidate drivers of patient outcome: both TFs have established oncogenic roles in ovarian cancer (Croft et al. 2024, Di Palma & Zannini 2022), but to our knowledge, neither TF’s copy-number-driven regulatory link to these specific target genes, nor its association with survival, has previously been reported, giving concrete, testable hypotheses for experimental follow-up.

The rest of the article is organized as follows. In Section 2, we introduce the empirical Bayes model underlying PRISM and describe the extensions to missing data, non-Gaussian modalities, grouped populations, and longitudinal data. In Section 3 we benchmark PRISM against existing methods on simulated data. In Section 4, we apply PRISM to a yeast cell-cycle time-course dataset, and in Section 5 we apply PRISM to paired copy number and expression data from TCGA ovarian cancer patients. Finally, we conclude with a discussion in Section 6. PRISM is distributed as part of the open-source Python package SIGMAnet.

## 2 Methods

### 2.1 Problem Setting

PRISM is motivated by cohorts in which each of *n* individuals contributes a high-dimensional molecular profile *x*_*i*_ ∈ ℝ^*g*^, and the scientific question is how the relationships among the *g* molecular features differ across individuals. These relationships are described by an individual-specific covariance matrix Σ_*i*_ (*g × g*), which is the object we wish to estimate for all *i* = 1, …, *n*. We illustrate this with two datasets.

#### Ovarian cancer

The individuals are *n* = 423 ovarian tumors. *x*_*i*_ denotes the stacked copy number of transcription factors (TFs) and the expression of genes for the *i*-th tumor, with *g* = 16,763. We estimate Σ_*i*_ and focus on the submatrix of Σ_*i*_ which record the copy number-expression covariances, where the entry linking TF *j* and gene *k* describes how, in tumor *i*, the dosage of the TF covaries with the expression of the gene. We call this a TF-gene edge which is a potential candidate marker of a regulatory mechanism active in that patient’s tumor. We examine whether these patient-specific edges are associated with survival outcome. Copy number data are non-Gaussian, which motivates the copula extension in Section 2.7.

#### Yeast cell cycle

The individuals are the *n* = 25 time points of a synchronized yeast cell cycle time course, where *x*_*i*_ contains the expression of *g* = 4,390 genes at time point *i*. The matrix Σ_*i*_ is the gene-gene coexpression at that time point, and we ask which gene pairs are coexpressed at which phase of the cell cycle. The temporal ordering motivates the longitudinal extension of PRISM described in Section 2.9.

In both applications we estimate one covariance matrix per individual from a single profile *x*_*i*_, which is possible only by borrowing strength across individuals. We start by developing PRISM for a generic *x*_*i*_, so that it applies to both. Sections 2.2-2.4 describes the core model, and Section 2.5 quantifies uncertainty for each edge. The subsequent sections extend the core model to features of real cohorts: non-Gaussian modalities such as copy number (Section 2.7, used for TCGA-OV), temporally ordered samples (Section 2.9, used for the yeast time course), and, although not used in these two applications, partially observed profiles and grouped populations, as in multi-omic cohorts with absent modalities, single-cell dropout, or subpopulations such as different ancestries (Sections 2.6 and 2.8).

### 2.2 Model Formulation

Let 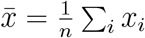 denote the sample mean. We assume that the centered observations satisfy

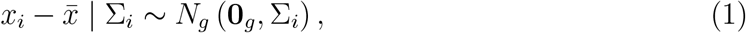

where Σ_*i*_ is the covariance matrix of the *i*-th observation. For a homogeneous population, Σ_*i*_ = Σ is the same for all individuals, but in most real data applications, where samples are not biological replicates, we expect Σ_*i*_ to vary across individuals.

We assume that Σ_*i*_ follows an inverse-Wishart prior distribution

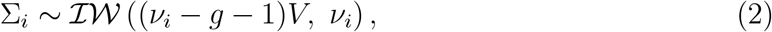

where *ν*_*i*_ *> g* + 1 denotes the prior degrees of freedom and *V* denotes the prior mean, shared across all individual covariances.

### 2.3 Estimating Prior Mean

Since 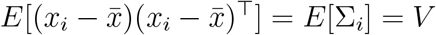, the sample covariance matrix

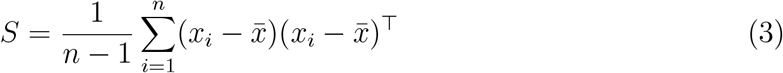

is an unbiased estimator of *V* for any *g, n* and *ν*_*i*_ *> g* + 1.

However, for *g ≥ n*, the sample covariance estimate *S* is ill-conditioned, which leads to the posterior mean of Σ_*i*_ being ill-conditioned as well. Many well-conditioned shrinkage estimators have been proposed as alternatives (Ledoit & Wolf 2004, Chen et al. 2010). So, for our software implementation we give the user the option to choose between *S* and the Oracle Approximating Shrinkage (OAS) estimator (Chen et al. 2010) as an estimate of *V*, as it has been shown to guarantee a well-conditioned, invertible estimate even when *g ≥ n*, while converging to *S* as *n → ∞* under Gaussian data. Our simulations show OAS matches *S*’s accuracy while remaining well-conditioned where *S* itself is singular.

### 2.4 Estimating Prior Degrees of Freedom

A natural approach is to estimate *ν*_*i*_ from the scaled Mahalanobis distance 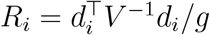 of each centered observation 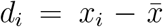. However, *E*[*R*_*i*_] = 1 for every value of *ν*_*i*_ (Supplementary Lemma 2), and the same holds for the pooled average 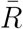. Hence any estimator that matches the mean of *R*_*i*_ to a function of *ν*_*i*_ is uninformative about *ν*_*i*_, however large *n* is. Information about *ν*_*i*_ lies instead in the variability of the individual covariances Σ_*i*_ around *V*, which motivates a variance-based calibration.

We therefore adopt the variance-matching calibration of Saha et al. (2024), which matches the prior variance of the diagonal entries of Σ_*i*_ under (2) to a jackknife estimate:

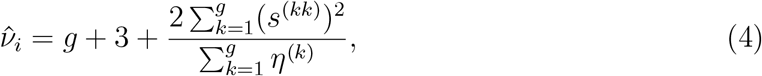

where *s*^(*kk*)^ is the *k*-th diagonal entry of *S* and *η*^(*k*)^ is the jackknife variance estimate for omics variable *k*. (4) guarantees 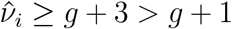 automatically, since the correction term is a ratio of a positive sum of squares to a nonnegative sum of variance estimates.

However, this calibration yields a common estimate 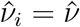 across individuals. If the data contain outliers, individual-specific estimates may be preferable. We leave this to future work.

Since Inverse-Wishart is a conjugate prior for the Gaussian covariance matrix, combining the Gaussian likelihood (1) with the Inverse-Wishart prior (2) gives the posterior

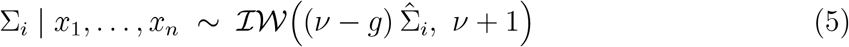

where

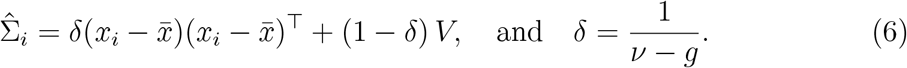

Since *ν ≥ g* + 1, we have 0 *< δ ≤* 1: small *δ* shrinks 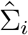 toward *V* (homogeneous populations), while large *δ* allows greater individual-specific deviations from the population-specific covariance estimate *V*. Our closest competing method BONOBO (Saha et al. 2024) let the prior mean itself vary by individual, recomputing a leave-one-out covariance *V*_*−i*_ for each individual, so that each individual is shrunk toward a prior mean that already excludes their own data. This leave-one-out construction departs from the standard empirical Bayes principle adopted here and also in SIREN (Saha 2026), where the prior is estimated once from the full population. This reduces the computational complexity of estimating *V* from *O*(*n*^2^*g*^2^) in BONOBO’s (Saha et al. 2024) leave-one-out approach to *O*(*ng*^2^) in ours.

### 2.5 Uncertainty Quantification for Individual Correlation Edges

We can derive posterior credible regions from the closed-form posterior 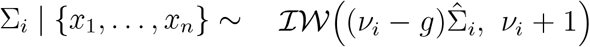. We provide two approaches: an exact Monte Carlo procedure requiring no distributional approximation, and a faster approach based on Normal approximation.

#### Exact posterior for a single edge

Any principal submatrix of an Inverse-Wishart-distributed matrix is itself Inverse Wishart distributed, with the degrees of freedom shifted by the difference in dimension (Mardia et al. 2024). Thus, for any pair of genes {*j, k*}, the exact joint posterior of the variance of the *j*-th gene, covariance of the *j*-th and *k*-th gene and the variance of the *k*-th gene, denoted by 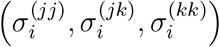, is

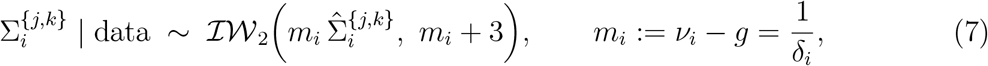

where 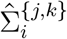 denotes the 2 *×* 2 principal submatrix of the posterior mean 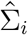 corresponding to genes *j* and *k*. Thus, we can obtain the exact posterior distribution of the correlation 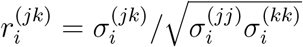 via direct Monte Carlo sampling.

From *B* posterior draws 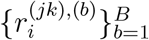, we construct equal-tailed credible intervals and a two-sided posterior *p*-value for the null hypothesis that the true coexpression entry is zero,

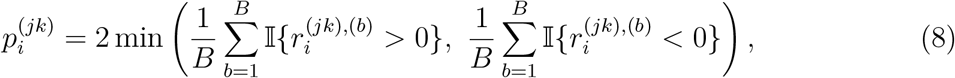

where I{*A*} denotes the indicator variable of an event *A*.

#### A fast closed-form alternative using normal approximation

Sampling Equation (7) for every edge requires repeated Monte Carlo draws which is computationally expensive. As a faster alternative, we approximate the posterior standard deviation of 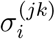 analytically,

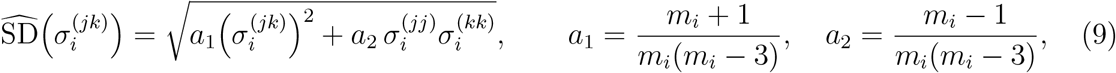

and construct a Wald-type interval and *z*-score *p*-value, 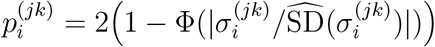. Unlike Equation (7), this is an approximation to the true marginal distribution of 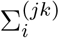. We validate the accuracy of this approximation against the exact method via simulation (Section 3.3) and demonstrate that it retains comparable and approximately nominal coverage to Monte Carlo sampling, under both the Gaussian and Gaussian copula (Section 2.7) extensions of PRISM, with the Normal approximation showing marginally better-calibrated coverage and Type I error at substantially lower computational cost.

The pseudo-count *m*_*i*_ used in the point estimate 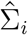 (Section 2.4) is calibrated by matching the prior variance of the diagonal entries of *S*, a criterion chosen to yield well-calibrated shrinkage of the posterior *mean*. This criterion does not itself guarantee that *m*_*i*_ correctly reflects how much is actually known about the population parameter *V*, which is estimated, not observed, from the *n* available samples. This is a general and well-documented limitation of empirical Bayes procedures: plugging in point estimates of hyperparameters and treating them as fixed for downstream inference fails to propagate the hyperparameters’ own estimation uncertainty, producing credible intervals that are too narrow, particularly when little data underlie the hyperparameter estimate (Morris 1983, Laird & Louis 1987). We find that the calibrated 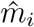 can substantially exceed *n*, implying via Equation (7) that *V* is known to a precision equivalent to far more independent observations than were actually collected. Since *V* cannot be known more precisely than the information content of the *n* samples used to estimate it, we impose a conservative cap,

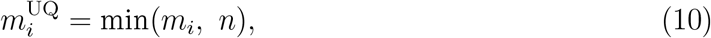

This cap applies only to uncertainty quantification and leaves the point estimate 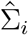 itself unaffected. We validate this calibration using simulated data (Section 3.3), confirming that the resulting 95% credible intervals achieve approximately nominal coverage.

### 2.6 Handling Missing Data

When analyzing multi-omic data, some individuals might have one omics modality entirely missing or both modalities partially missing. The problem of missing data is even more prevalent for single-cell omics data, which are frequently affected by dropout, in which a subset of variables is unobserved for a given sample. Suppose that for sample *i*, the coordinates of *x*_*i*_ partition into an observed set *O*_*i*_ and a missing set *M*_*i*_. Because marginals of a multivariate Normal are themselves Normal, the likelihood contribution of sample *i* depends only on the observed sub-vector 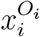 and the corresponding sub-matrix 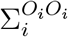; the missing coordinates contribute no information to the likelihood. Applying the same conjugate update as above to this reduced likelihood shows that the posterior mean of the observed block of 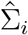 follows (6) as before. The posterior mean of the blocks involving *M*_*i*_ also has a closed form (Supplementary Lemma 3), but evaluating it requires solving a linear system in the observed block 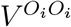 for every sample, which costs up to *O*(*g*^3^) per distinct missingness pattern and does not scale to the dimensions and sample sizes of omics data. For computational efficiency, we therefore approximate every entry involving at least one variable in *M*_*i*_ by the corresponding entry of the prior mean *V* :

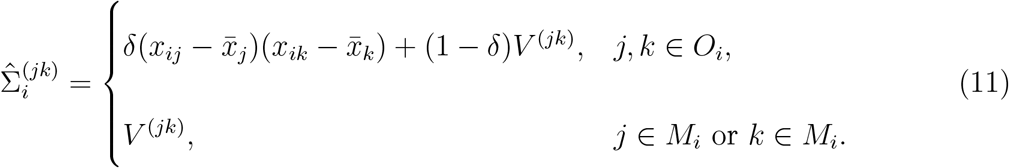

This approximation ignores the information that the observed data carry about the entries involving *M*_*i*_ through their dependence on the observed block, and the resulting error is of order *δ* = 1*/*(*ν − g*), and therefore small when the prior is concentrated around *V*.

We note that this update is based on the assumption of missing-completely-at-random, which might be too strong for some single-cell data. The mechanism underlying zero counts in single-cell data remains debated, with some zeros reflecting true absence of expression and others reflecting technical undersampling, but in either case disproportionately affecting genes with lower expression levels (Hicks et al. 2018). Extending the closed-form update in Equation (11) to explicitly model a missing-not-at-random mechanism might involve incorporating prior knowledge about the missingness structure into the data likelihood itself, for example through a zero-inflated hurdle-type model (McDavid et al. 2019, Cui & Wang 2023) where the probability of missingness depends on the latent true expression level of the gene or a Poisson measurement model (Sarkar & Stephens 2021) where the probability distribution of the measured expression depends on the true unobserved expression level of a gene. Such an extension would likely forgo the closed-form conjugate update available under the simpler missing-completely-at-random assumption, and we leave it to future work.

### 2.7 Extending for Non-Gaussian Omics Data

The Gaussian assumption in (1) is reasonable for continuous unimodal omics data, such as normalized microarray data or RNA sequencing data after log-transformation, but can be violated for other omics modalities, such as DNA methylation or Copy Number Alteration (CNA) data. We relax this assumption using a Gaussian copula model (Liu et al. 2009), where we assume that there exist unknown monotone increasing transformations *f*_1_, …, *f*_*g*_ such that

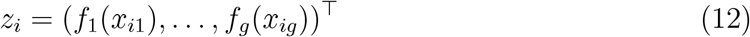

follows the multivariate Normal model in (1). We estimate *f*_*j*_ empirically for each omics variable *j* via a marginal Gaussianizing transform based on the empirical distribution of *x*_*·j*_ across samples, and apply the posterior estimation framework described above to the transformed data *z*_*i*_ in place of *x*_*i*_: *V* and the individual-sample posterior mean term are both computed directly from *z*, exactly as in Section 2 for Gaussian data.

Because Cov(*X*)*≠* Cov(*Z*) in general under a nonlinear monotone transformation, PRISM’s edges reflect association on the latent scale *Z* and not on the original data *X*. The sign of each association is preserved but the magnitudes are not. This is sufficient for many practical purposes: identifying candidate regulatory interactions for clinical intervention or experimental follow-up typically requires knowing the *direction* of an association, e.g., whether increased dosage of a regulator is associated with increased or decreased target expression, rather than its exact magnitude on the original scale *X*. A fully hierarchical model jointly estimating *f*_1_, …, *f*_*g*_ and Cov(*X*) directly would avoid this scale mismatch altogether, but is computationally prohibitive at the dimension of omics data.

### 2.8 Extending for Grouped Data

PRISM’s pooled prior *V*, estimated once from the full sample, is designed for settings with mild, continuous inter-individual heterogeneity, which is the common case in omics cohorts where individuals do not fall along a known discrete structural divide. When individuals instead fall into severely distinct, structurally-driven subpopulations, for example, samples drawn from distinct tissues, pooling all samples into a single *V* is inappropriate, since *V* would not represent either subpopulation well. When group membership is known and reflects such severe, structural heterogeneity, we recommend estimating a separate prior mean *V*_*k*_ for each group *k*, using only that group’s samples, and applying the PRISM estimator independently within each group. This requires no modification to the underlying model or estimators. It is a direct application of the existing framework to a group-partitioned dataset rather than the full cohort.

When group labels are unavailable but the underlying population is known to consist of strong, discrete subpopulations, an ad-hoc approach would be to perform unsupervised clustering through conventional models such as k-means, Louvain or Leiden clustering (Steinley 2006, Traag et al. 2019) followed by group-specific PRISM estimation within each discovered cluster. We caution that clustering on the same data subsequently used to estimate *V*_*k*_ can understate uncertainty in the group assignments themselves, and recommend this approach only when subpopulation structure is visually or statistically unambiguous, rather than as routine preprocessing. As future work, we will explore combining mixture-models with PRISM as a hierarchical structure to automatically discover underlying clusters if present and propagate the uncertainty in cluster estimation to the posterior uncertainty of individual-specific covariance estimation. This discussion addresses severe, structural heterogeneity specifically and is not intended for the milder, continuous heterogeneity PRISM’s pooled-prior design already accommodates via shrinkage (Section 3.1). We note that unnecessary group-splitting in the absence of severe structural differences would only reduce the effective sample size available for estimating each *V*_*k*_, likely affecting accuracy.

### 2.9 Extending for Longitudinal Omics Data

Many omics studies profile the same biological system, a cell population, a developing organism, a patient under treatment at a sequence of time points rather than as a single cross-section. A common ad hoc practice for such data is a sliding/rolling-window analysis (Naoki et al. 2025, Karwowska et al. 2024): computing a summary statistic (e.g., a correlation or covariance matrix) separately within each of several overlapping windows of nearby time points, without a formal probabilistic model connecting the windows or quantifying uncertainty in the resulting estimates. We show that PRISM’s hierarchical framework accommodates this setting directly, replacing the ad hoc rolling window with a local empirical Bayes prior, providing closed-form posterior estimation at every time point.

Suppose the *n* samples are ordered by time (or pseudotime (Trapnell et al. 2014)), indexed *i* = 1, …, *n* according to this ordering. For a chosen window with some integer half-width *k*, define the local neighborhood of time point *i* as

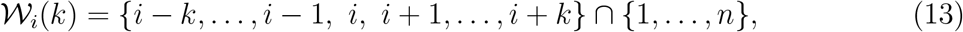

truncated at the boundaries of the series when *i ≤ k* or *i > n − k*, so that early and late time points use an asymmetric, smaller window of available neighbors rather than being excluded. In place of the single population-level prior mean *V* of Section 2.3, we estimate a local prior mean 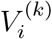 from the samples in *W*_*i*_(*k*),

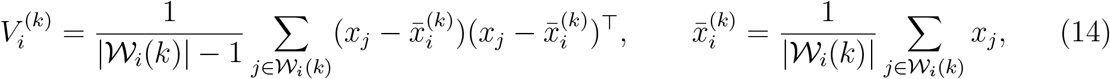

and similarly recalibrate the prior degrees of freedom *ν*_*i*_ (equivalently *δ*_*i*_) via the jackknife estimator of Equation (4), applied to the same local window *W*_*i*_(*k*) rather than to the full data. The conjugate posterior update is then applied at each time point *i*, using 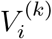 in place of *V* and the locally-calibrated *δ*_*i*_, yielding a posterior mean 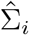, and edge-level credible intervals. The Gaussian copula extension of Section 2.7 applies analogously: the marginal Gaussianizing transform *f*_*j*_ is estimated from the samples in *W*_*i*_(*k*) rather than from the full cohort, since a transform fit globally across all time points may not correctly Gaussianize the local marginal distribution at a given time if the marginals themselves vary over time.

This construction is built on the assumption that the underlying coexpression structure varies smoothly over time, so that nearby time points are informative about a given time point’s own structure, an instance of *local stationarity* (Dahlhaus 1997, Chen, Li, Li & Linton 2025). This assumption can be violated at sharp regulatory transitions, such as cell-cycle phase changes where coexpression structure may shift over a short time interval. Even in such cases, however, discretely sampled time points rarely capture a truly instantaneous transition, as genes and pathways within a regulatory program typically transition asynchronously rather than simultaneously, unfolding as a sequence of staggered waves rather than a single synchronized switch (Trapnell et al. 2014), so that some degree of local smoothness is usually preserved at the resolution of realistic sampling. The smaller the window *k*, the less this construction relies on smoothness holding over any particular interval.

The window size *k* governs bias-variance tradeoff: larger *k* incorporates more samples into 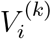, reducing its estimation variance, but risks averaging over genuine time-variation (bias) if the underlying structure changes considerably within the window, whereas smaller *k* captures local structure more faithfully but yields a noisier, and at small *k* potentially ill-conditioned local prior, particularly when *g* is large relative to the local window size 2*k* + 1. In such situations we recommend using the OAS prior (Chen et al. 2010).

#### 2.9.1 Selecting *k* through cross-validation

To select *k* through cross-validation, we compute the local prior 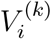 from *W*_*i*_(*k*), excluding sample *i* itself, and evaluate the predictive log-likelihood of the held-out sample *x*_*i*_ under the resulting local marginal-*t* model. Averaging this predictive score over all time points gives a cross-validated criterion as a function of *k*. We select the value of *k* that maximizes it. This is analogous to standard bandwidth-selection procedures for local likelihood and local polynomial estimation (Fan 2018, Ruppert et al. 1995), and more broadly, to bandwidth selection in locally stationary time series (Dahlhaus 1997). Related principles for choosing a smoothing bandwidth, such as autocorrelation-function (ACF)-decay analysis, frequency-domain filtering (Dahlhaus 1997), and analytical mean-squared-forecast-error optimization (Ruppert et al. 1995) are largely equivalent formulations of the same underlying bias-variance tradeoff, and in principle any of these approaches could be adapted here.

#### 2.9.2 Selecting *k* through split-half structural consistency

Leave-one-out predictive log-likelihood favors whichever window most improves overall covariance estimation accuracy and hence is dominated by the sheer number of well-estimated off-diagonal entries, rather than by the subset of entries that actually vary over time, which might be much smaller in general. Consequently, when the true structure changes only within a limited window around each time point, the log-likelihood objective can still favor a larger window, because pooling more samples improves overall predictive accuracy by more than the cost of blurring that local change. As a result, the log-likelihood criterion tends to select *k* toward larger values, even when a smaller window would better capture the time-localized structure.

Therefore, we additionally implement split-half structural consistency which is designed to be directly sensitive to local homogeneity rather than to overall estimation accuracy. For each time point *i* and candidate *k*, we partition the window into a “left” half consisting of the samples preceding *i* within *W*_*i*_(*k*) and a “right” half consisting of the samples following *i*. A local covariance estimate is computed independently on each half using the OAS estimator. The Pearson correlation between the off-diagonal entries of the two half-estimates serves as a measure of structural agreement at that time point and window width. We compute a split-half consistency score for each *k* by averaging over all time points for which both halves contain at least *h*_min_ samples (`min_half_size` in the software with default choice 15), and select *k* that maximizes this average score. Time points for which a half falls below *h*_min_ are excluded from that candidate *k*’s score. Split-half consistency is also considerably cheaper to compute than the leave-one-out log-likelihood search.

This criterion is an instance of a more general principle of selecting a local window or bandwidth by comparing sub-interval estimates for mutual agreement, named *local change point* (LCP) analysis (Mercurio & Spokoiny 2004, Spokoiny 2009), where the interval of local homogeneity for a (univariate) time-varying parameter is recovered by testing whether estimates from different sub-intervals agree. We adapt this principle to the multivariate setting of sample-specific covariance estimation. In place of the sequential hypothesis-testing procedures typically used in the univariate LCP literature, we use a simple Pearson-correlation agreement score between the OAS covariance estimates from the two half-windows.

We select a single window size *k* shared across all genes, rather than allowing windows tailored to each variable’s autocorrelation. This simplification is deliberate to preserve positive semi-definiteness of the prior as assembling a local covariance entrywise from gene-specific windows is not guaranteed to be positive-semi-definite. A less restrictive extension, which we leave to future work, would allow the window size *k*_*i*_ to vary smoothly with time (but not with gene), continuing to use a single shared window across all genes at each time point while adapting its width locally to the estimated rate of change of the joint correlation structure.

## 3 Applications on Simulated Data

The simulations are designed to mirror the features of the two applications. TCGA-OV tumors are heterogeneous and their copy number data are non-Gaussian, so we vary the degree of heterogeneity between individuals (Simulation 1) and use non-Gaussian marginals (Simulation 2). Because the TF-gene edges identified in TCGA-OV are interpreted individually, we assess the calibration of edge-level uncertainty (Simulation 3), and its behavior when some entries are missing (Simulation 4). The yeast time course has samples that are ordered in time, so we evaluate the longitudinal extension on series with smooth and abrupt changes (Simulation 5). Both applications have far more genes than samples, so the simulations include regimes with *g ≥ n*.

We evaluate PRISM against BONOBO (Saha et al. 2024) and SWEET (Chen et al. 2023) on simulated data with a known, sparse true correlation structure, repeating each experiment across 5 replicates. Although PRISM allows each individual’s covariance Σ_*i*_ to deviate from the population-level *V*, it does not presuppose that the population consists of discrete, heterogeneous subgroups. We therefore benchmark on simulated data from both homogeneous and heterogeneous (mixture) populations, to confirm that PRISM performs well under both, since it is often impractical to know in advance whether the population underlying a given omics dataset is homogeneous or heterogeneous.

For homogeneous case, to construct the true correlation matrix, we start with the identity matrix and assign a proportion of off-diagonal entries to nonzero values drawn uniformly from [*−*1, 1]. The diagonal entries are then set to the row-sum of absolute off-diagonal entries plus a small positive value *ϵ* = 0.0001, and the matrix is rescaled to unit diagonal.

Thus, the resulting matrix becomes diagonally dominant and hence positive definite. As dimension *g* varies, the proportion of active off-diagonal entries is scaled as *g* grows so that the expected number of true edges per gene remains fixed (target degree = 5).

Heterogeneous populations are simulated as a two-group mixture: the first group’s correlation structure is simulated as above. The second group’s true correlation matrix is constructed by removing a fraction *p* of the first group’s true edges and adding an equal fraction *p* of new edges at previously-zero positions, then reapplying the same diagonal-dominance construction to the perturbed edge set. Thus, *p* = 0 corresponds to a homogeneous population and larger *p* corresponds to greater heterogeneity between the two groups.

For each sample, we compute the mean squared error (MSE) and Pearson correlation between the estimated and true correlation matrices, and the area under the ROC curve (AUC) for recovering the true edge set. We report the mean of each metric across samples.

### 3.1 Simulation 1: Gaussian Data

In *simulation setting 1*, for each simulated dataset, we draw the individual omics profiles from a multivariate Gaussian distribution with the corresponding group’s true correlation matrix, with independent Gaussian measurement noise (sd = 0.1) added to each entry. We compare PRISM, using both the sample covariance (denoted PRISM(sample)) and the OAS estimator (Chen et al. 2010) (denoted PRISM(OAS)) as the prior mean *V*, against BONOBO (Saha et al. 2024) and SWEET (Chen et al. 2023). We compare performance of the four methods for varying sample size *n* ∈ {50, 100, 500, 1000}, keeping fixed *g* = 500 and varying dimension *g* ∈ {50, 100, 500, 1000}, keeping fixed *n* = 500). In both experiments, we keep the heterogeneity proportion fixed at *p* = 0.1. Then we vary heterogeneity *p* ∈ {0, 0.1, 0.3} keeping *n* = 100 and *g* = 500 fixed.

We observed (Figure 2 (a)-(c)) that PRISM (sample), PRISM (OAS), and BONOBO achieve nearly identical AUC throughout, indicating that PRISM’s estimator based on a common prior distribution across all samples recovers the true edge structure as well as BONOBO, which assumes that the prior mean and degrees of freedom parameters vary across individuals. SWEET consistently underperforms across all experiments. AUC drops sharply from the homogeneous case (*p* = 0) to any nonzero heterogeneity and then plateaus, for every method considered. This indicates that even mild heterogeneity substantially degrades edge recovery in general, rather than being a limitation specific to any one estimator.

**Figure 2:**
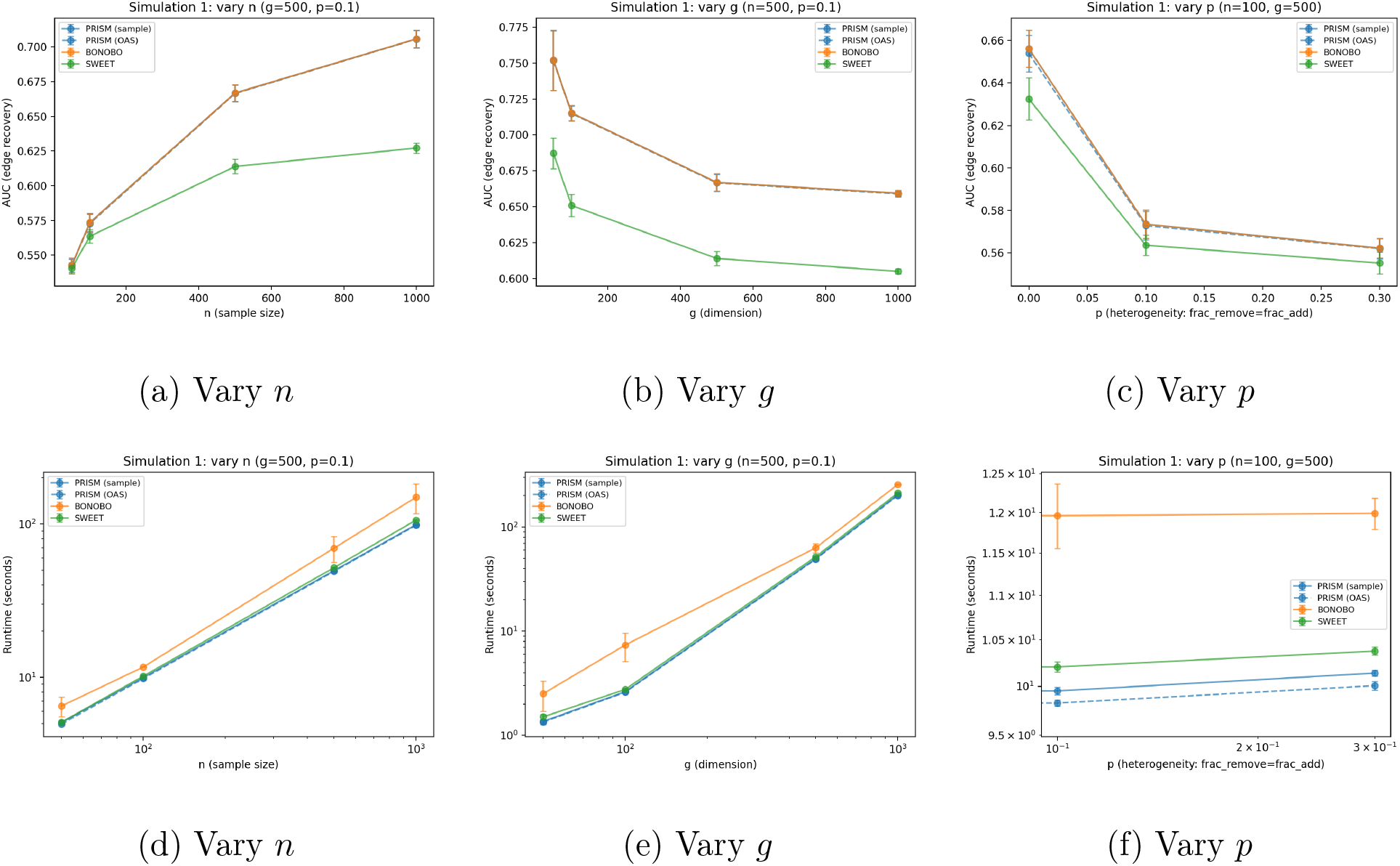
Simulation 1 (PRISM vs. BONOBO vs. SWEET): edge-recovery AUC (top row) and runtime (bottom row) as a function of sample size *n*, dimension *g*, and heterogeneity *p* (fraction of true edges removed/added between groups).

PRISM and SWEET have comparable runtime (Figure 2 (d)-(f)), while BONOBO is consistently slower, consistent with BONOBO’s *O*(*n*^2^*g*^2^) cost relative to PRISM’s *O*(*ng*^2^) cost.

While PRISM’s MSE is substantially lower than BONOBO’s throughout (Supplementary Figure S1), PRISM’s Pearson correlation with the truth closely coincides with that of BONOBO’s. Since Pearson correlation does not show a corresponding PRISM advantage, the MSE gap reflects PRISM’s shrinkage producing estimates of smaller overall magnitude rather than a proportionally better recovery of the underlying correlation structure.

### 3.2 Simulation 2: Non-Gaussian Data

We repeat the same experiments (varying *n, g* and *p*) as in simulation setting 1, on data with non-Gaussian marginals, generated by applying a Beta(0.2, 0.2) transform (mimicking DNA methylation beta-values) to the latent Gaussian draws underlying the same sparse heterogeneous ground truth as Simulation 1. We compare standard PRISM (sample covariance and OAS priors) and the Gaussian copula extension of PRISM (PrismMultiomic in figures; with sample and OAS priors), with BONOBO and SWEET, all evaluated against the true *latent* correlation structure.

The Gaussian copula PRISM variants perform comparably or slightly better than the standard PRISM and BONOBO in AUC (Figure 3 (a)-(c)). SWEET underperforms all other methods. AUC declines sharply from homogeneous case to any nonzero heterogeneity and then plateaus. MSE and Pearson correlation follow similar trend (Supplementary Figure S2).

**Figure 3:**
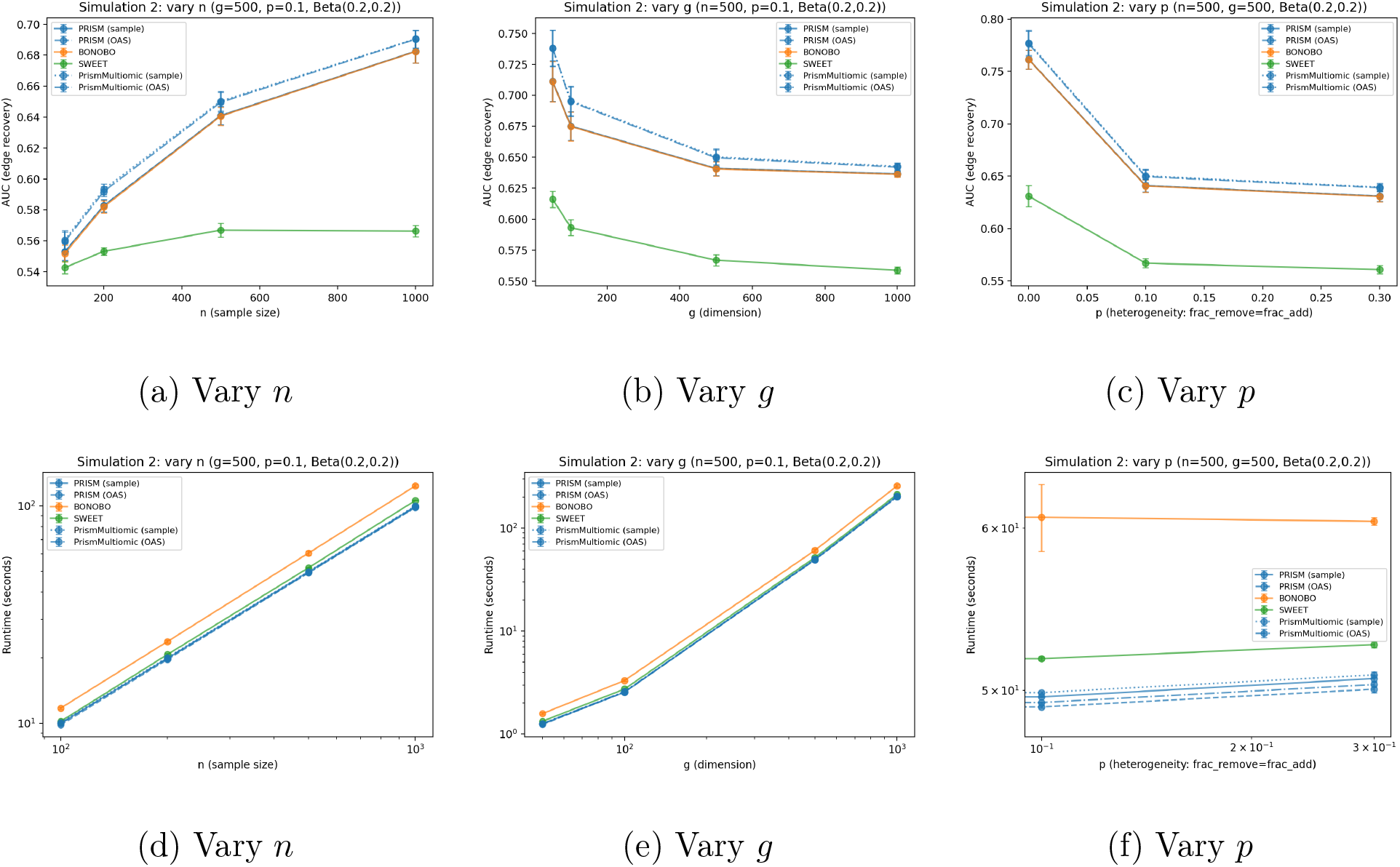
Simulation 2 (PRISM vs. Gaussian Copula PRISM vs. BONOBO vs. SWEET, Beta(0.2,0.2)-transformed data): edge-recovery AUC (top row) and runtime (bottom row) as a function of sample size *n*, dimension *g*, and heterogeneity *p*.

In runtime (Figure 3 (d)-(f)), BONOBO remains the slowest method throughout. The copula PRISM variant has a slightly higher computation cost over standard PRISM due to the marginal-transform estimation step, but remain substantially faster than BONOBO.

We provide a supplementary analysis (Supplementary Figure S3) to check robustness of the Gaussianizing transformation under alternative distributions, using Negative Binomial marginals (mimicking CNA read-depth data) in place of the Beta transform considered in Simulation 2. Under the tie-heavy Negative Binomial marginals, all PRISM variants (standard and Gaussian copula) achieve comparable AUC, indicating no detectable benefit of the Gaussianizing transformation in this setting, consistent with rank-based Gaussianizing transformation being less effective when many observations share identical values.

Our simulation experiments show that applying the copula transformation to data that is already approximately Gaussian introduces a negligible accuracy cost, alongside a modest increase in computation time from fitting a marginal transform that is unnecessary when the true marginals are already Gaussian. We therefore recommend checking marginal normality (e.g., via a Shapiro-Wilk test) before applying the Gaussian copula variant of PRISM, defaulting to standard PRISM when marginals are already approximately Gaussian.

### 3.3 Simulation 3: Uncertainty Quantification

We evaluate edge-level posterior uncertainty calibration via three simulation studies and report the median and interquartile range of per-sample coverage, pooled across all evaluated individuals and simulation replicates.

We first assess whether the fast, closed-form *z*-score approximation (Equation (9)) achieves coverage and Type I error comparable to the exact posterior (Equation (7)), which we evaluate using two implementations described in Section 2.5: a per-pair Monte Carlo loop and a vectorized batch sampler that target the same posterior and, as expected, give closely agreeing coverage and Type I error throughout. We recommend the vectorized implementation in practice for its substantially lower computational cost (Supplementary Figure S4).

We additionally examine whether PRISM’s posterior maintains approximately nominal coverage as dimension increases, comparing *g* = 50 against *g* = 200 under both homogeneous and heterogeneous (two-group mixture) populations, with sample size fixed at *n* = 100 throughout so that the two dimensions considered span both the *n > g* and *n ≤ g* regimes.

For each setting, we compute 95% credible intervals and two-sided *p*-values for every edge of every evaluated individual across 5 simulated replicates, using both the exact and considerably faster *z*-score approaches, and report empirical coverage and Type I error (Supplementary Table S1). Both approaches achieve approximately nominal coverage at *g* = 50 and *g* = 200 under the homogeneous population, with modestly reduced coverage under heterogeneity.

At *g* = 50, we additionally compare PRISM’s coverage (here, the *z*-score approximation) against the edge-level uncertainty derived from BONOBO and SWEET, on the same Gaussian ground truth as Simulation 1. We use BONOBO’s own native significance-testing mechanism for comparison. SWEET has no native probabilistic model, so we construct a nonparametric bootstrap proxy for its uncertainty, resampling the individuals contributing to its population-level correlation estimate. Because this reflects resampling variability of a point estimate rather than a Bayesian posterior, it is not a fully like-for-like comparison and is interpreted accordingly. Given the substantially higher computational cost of BONOBO’s leave-one-out recomputation and SWEET’s bootstrap relative to PRISM’s closed-form posterior, we restrict the comparison to *g* = 50, evaluating all individuals for all three methods. PRISM’s own coverage was separately confirmed to remain approximately nominal at *g* = 200 (above), suggesting this restriction is unlikely to qualitatively change the comparison’s conclusions. PRISM achieves coverage close to nominal in both the homogeneous and heterogeneous settings, comparable to BONOBO while SWEET under-performs.

We repeat this three-method comparison on the Beta(0.2, 0.2)-transformed (DNA methylation-like) ground truth of Simulation 2 (Table 2), evaluating both PRISM’s exact and *z*-score uncertainty quantification variants (PrismMultiomic in figures) alongside BONOBO and SWEET. Both PRISM variants retain approximately nominal coverage and well-controlled Type I error under the transform, in both populations. BONOBO’s native uncertainty quantification, by contrast, coverage falls to approximately 0.14 with Type I error near 0.84, indicating that its Normal-approximation significance test, calibrated under an implicit Gaussian likelihood, is not robust to non-Gaussian marginals, even though its point estimates (Section 3.2) are computed on the same untransformed data without similar performance drop. SWEET’s bootstrap proxy remains under-covered as in the Gaussian example.

**Table 1:** Uncertainty quantification comparison on Gaussian data: PRISM (z-score) vs. BONOBO’s native uncertainty quantification vs. SWEET’s bootstrap proxy, at *g* = 50. Coverage and Type I error are median values pooled across all evaluated individuals and 5 simulation replicates (*n* = 100); IQR is the interquartile range of per-sample coverage. Nominal targets: coverage = 0.95, Type I error = 0.05.

| Population | Method | Coverage | IQR | Type I |
| --- | --- | --- | --- | --- |
| Homogeneous | PRISM | 0.961 | 0.955-0.961 | 0.041 |
|  | BONOBO | 0.946 | 0.936-0.951 | 0.053 |
|  | SWEET | 0.859 | 0.835-0.885 | 0.130 |
| Mixture (2 groups) | PRISM | 0.937 | 0.936-0.941 | 0.045 |
|  | BONOBO | 0.940 | 0.936-0.950 | 0.040 |
|  | SWEET | 0.851 | 0.826-0.869 | 0.122 |

**Table 2:** Uncertainty quantification comparison on Beta(0.2,0.2)-transformed data: PRISM’s exact and z-score variants vs. BONOBO vs. SWEET, at *g* = 50. Coverage and Type I error are median values pooled across all evaluated individuals and 5 simulation replicates (*n* = 100); IQR is the interquartile range of per-sample coverage. Nominal targets: coverage = 0.95, Type I error = 0.05.

| Population | Method | Coverage | IQR | Type I |
| --- | --- | --- | --- | --- |
| Homogeneous | PRISM (exact) | 0.940 | 0.936-0.944 | 0.056 |
|  | PRISM (z-score) | 0.947 | 0.945-0.953 | 0.046 |
|  | BONOBO | 0.138 | 0.130-0.145 | 0.850 |
|  | SWEET | 0.858 | 0.829-0.882 | 0.128 |
| Mixture (2 groups) | PRISM (exact) | 0.922 | 0.919-0.925 | 0.060 |
|  | PRISM (z-score) | 0.944 | 0.943-0.949 | 0.048 |
|  | BONOBO | 0.146 | 0.122-0.154 | 0.834 |
|  | SWEET | 0.851 | 0.827-0.873 | 0.117 |

### 3.4 Simulation 4: Missing Data Handling

We assess PRISM’s missing-data handling capabilities on the same sparse heterogeneous ground truth as above, masking a proportion of entries from the expression data independently at each cell, under a missing-completely-at-random (MCAR) mechanism. For each simulated individual, we distinguish *observed edges* (both genes present for that individual) from *fallback edges* (at least one gene missing, so the posterior mean falls back exactly to the prior), and evaluate PRISM’s *z*-score edge-level uncertainty quantification at *g* = 50 and *g* = 200 (*n* = 100, 5 replicates, 10% missingness).

At *g* = 50, median per-sample coverage (Supplementary Figure S5) is comparable for observed (0.914) and fallback (0.911) edges, indicating that the fallback to prior does not introduce edge-type-specific miscalibration beyond the modest under-coverage already present in this heterogeneous setting (Section 3.3). Coverage and Type I error rates at both dimensions, for missingness levels ranging from 10-50%, are reported in Supplementary Figure S6, showing performance declining with increased missingness, as expected.

### 3.5 Simulation 5: Time-Varying Data

We evaluate the longitudinal extension of PRISM (`PrismLongitudinal`) against two base-lines: (i) standard *naive PRISM*, the same estimator fit once using all time points (ignoring temporal order entirely), and (ii) a *rolling-window* baseline, the plain sample Pearson correlation matrix computed within the same local window of size 2*k* + 1 used by `PrismLongitudinal`, which intends to assess whether PRISM’s empirical-Bayes shrinkage improves recovery beyond windowing alone.

We simulate a single time-ordered series of gene expression over *n* = 100 time points and *g* = 200 genes, under two regimes governing how the true correlation structure evolves over time: (a) **linear drift**, in which the true correlation matrix linearly interpolates between two independently generated sparse endpoint matrices across the series and (b) **stepwise drift**, in which the series is partitioned into 4 contiguous, equal-length regimes (change points at *t ≈* 25, 50, 75), each assigned an independently generated true correlation matrix, with an abrupt change at each boundary. Both regimes follow the diagonally-dominant construction and noise model described before. Each experiment is repeated across 5 replicates.

#### Window selection

For linear drift, log-likelihood cross-validation selected *k* = 49 (the largest candidate considered, spanning nearly the full series) at a one-time cost of 52.9s. For stepwise drift, log-likelihood CV again selected a large window (*k* = 47, cost 48.1s), despite the presence of 3 genuine change points. By contrast, the split-half structural consistency criterion selected *k* = 15 for stepwise drift, at a one-time cost of only 1.8s.

To assess which selection better serves edge recovery, we additionally varied *k* and compared against the known ground truth on the stepwise dataset (Supplementary Table S2). AUC and Pearson correlation both show a clear interior optimum, peaking at *k* = 20 (AUC= 0.598) and declining toward both smaller *k* (insufficient samples for stable local estimation) and larger *k* (blurring across regime boundaries), including at the log-likelihood-selected *k* = 47 (AUC= 0.557). Split-half’s selected *k* = 15 achieves AUC= 0.594, within 0.004 of this oracle optimum and 0.037 above log-likelihood CV, despite using no ground-truth information. We therefore use log-likelihood CV’s selected *k* for linear drift and split-half’s selected *k* for stepwise drift as the window widths used by PrismLongitudinal and the rolling-window baseline in the comparisons below. We report MSE, Pearson correlation, AUC, and per-fit runtime across 5 replicates for both drift regimes (Table 3).

**Table 3:** Results for longitudinal data: AUC and per-fit runtime, mean *±* SD across 5 replicates, *g* = 200, *n* = 100.

| Drift | Method | AUC | Runtime (s) |
| --- | --- | --- | --- |
| Linear | **PrismLongitudinal** | $0.603 \pm 0.008$ | $0.123 \pm 0.025$ |
| | Naive PRISM | $0.625 \pm 0.009$ | $0.035 \pm 0.003$ |
| | Rolling window | $0.604 \pm 0.008$ | $0.463 \pm 0.003$ |
| Stepwise | **PrismLongitudinal** | $0.589 \pm 0.006$ | $0.093 \pm 0.007$ |
| | Naive PRISM | $0.539 \pm 0.005$ | $0.036 \pm 0.006$ |
| | Rolling window | $0.594 \pm 0.008$ | $0.162 \pm 0.002$ |

For the *linear drift* example, all three methods achieve comparable Pearson correlation with the truth (Supplementary Table S3). `PrismLongitudinal` and the rolling-window baseline achieve comparable AUC (Table 3) while naive PRISM achieves somewhat higher AUC, as no individual time point’s true structure differs sharply from the population average.

For the *stepwise drift* example `PrismLongitudinal` and the rolling-window baseline again achieve comparable Pearson correlation and AUC, both now substantially exceeding naive PRISM, demonstrating that once a genuine regime change is present, local windowing meaningfully improves recovery of the true correlation structure over a globally pooled fit.

Given a fixed, already-selected *k*, `PrismLongitudinal` is slower than naive PRISM (Table 3) but faster than the rolling-window baseline, and, unlike the rolling-window baseline, additionally provides calibrated per-edge credible intervals and *p*-values at every time point. The one-time cost of window selection is substantially larger than any single fit under log-likelihood CV (*∼* 48-53s) but negligible under split-half consistency (1.8s).

Together, these results indicate that `PrismLongitudinal’s` benefit over naive pooling is asymmetric: comparable accuracy at modest additional cost under smooth drift, and a substantial accuracy advantage under abrupt drift, while consistently matching the rolling-window baseline’s edge-recovery accuracy at lower runtime and with the added benefit of calibrated uncertainty quantification. Thus `PrismLongitudinal` is the safer default in real data situations where the smoothness of the true underlying process is unknown.

## 4 Application on Yeast Cell-Cycle Data

The goal of this analysis is to test whether PRISM recovers time-varying coexpression structure in a system whose underlying biology is well characterized. We applied PRISM to a cell-cycle time-course gene expression dataset from budding yeast (Pramila et al. 2006) (Downloaded from Gene Expression Omnibus with accession number GSE4987). First, we assessed the robustness of PRISM network estimates through reproducibility of coexpression networks between two replicates. Then we established the biological relevance of PRISM networks by demonstrating that they recover cell-cycle biology independently documented in existing literature.

W303a yeast cells were arrested with alpha factor in YEPD media. After release from arrest, the cells were sampled at 5-minute intervals for two hours spanning two full cell cycles, in two independent biological replicates, generating two microarray datasets. We downloaded gene expression values provided as Rosetta Resolver-normalized two-color log-ratios from the Gene Expression Omnibus (Edgar et al. 2002). We removed genes with missing values in at least one replicate, and further filtered out genes with low variance, retaining only 75% of genes with the largest minimum variance across the two replicates. This yielded an identical set of 4,390 genes in both replicates, which were then z-score normalized independently within each replicate across its own time points.

We first verified whether the window size *k* for PRISM longitudinal is consistent across the two independent replicates. Applying the split-half consistency selection criterion to each replicate separately, without reference to the other, both replicates independently selected *k* = 6, showing that PRISM infers the same local temporal neighborhood size, and hence the same effective time scale over which the coexpression structure is locally homogeneous from two independently synchronized biological cultures. Because each replicate has only 25 time points, we lowered *h*_min_ from its default of 15 to 1 and searched *k ∈* 1, …, 24.

Next, we assessed the reproducibility of PRISM networks at the level of individual coexpression edges. For each of the approximately 9.6 million 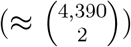 gene pairs, we computed the Pearson correlation between the two replicates’ full time-resolved coexpression networks. The aggregate median correlation across all edges was *r* = 0.487. However, since only a minority of gene pairs are expected to show genuine cell-cycle-linked coexpression, we stratified edges into deciles by within-replicate signal strength (variance of the coexpression trace). Reproducibility increased monotonically with signal strength, from a median *r* = 0.13 in the lowest-variance decile to *r* = 0.88 in the highest-variance decile (Supplementary Figure S9). This demonstrates that reproducibility of time-varying coexpression networks across the two replicates scales monotonically with signal-to-noise ratio, rather than reflecting a sharp division between reproducible and non-reproducible edges.

To verify whether known cell-cycle-specific gene–gene interactions are captured by PRISM, we examined the time-varying coexpression trajectories of six gene pairs whose regulatory relationships are well established in the yeast cell cycle literature (Figure 4). SWI5, a TF, enters the nucleus during the late M phase and triggers a sharp increase in transcription of *SIC1*, and directly activates *EGT2* at the end of mitosis and in early G1 (SWI5-EGT2). Both edges show pronounced cyclical coexpression with peaks at closely matching time points (approximately 35 and 90–95 minutes), consistent with both genes being targets of the same TF within the same phase window (Cherry et al. 1998). In contrast, SWI5 and its paralog ACE2 are co regulated across the cell cycle by similar upstream inputs rather than one directly activating the other (Schreiber et al. 2025, Wu et al. 2016). Accordingly, SWI5-ACE2 coexpression remains positive throughout the time course, even as its magnitude fluctuates. Similarly, FKH1, together with its paralog FKH2, coordinates the transcriptional timing of the G2/M cyclins *CLB1* and *CLB2* (FKH1-CLB1, FKH1-CLB2) (Kumar et al. 2000, Pic-Taylor et al. 2004). Both edges peak at approximately 25–30 minutes, earlier than the SWI5-SIC1 and SWI5-EGT2 peaks, consistent with the the fact that the G2/M cluster is transcribed prior to the M/G1 cluster in cell cycle. FKH1 and FKH2 themselves remain positively coexpressed throughout, as expected for joint activators of this program (FKH1-FKH2).

**Figure 4:**
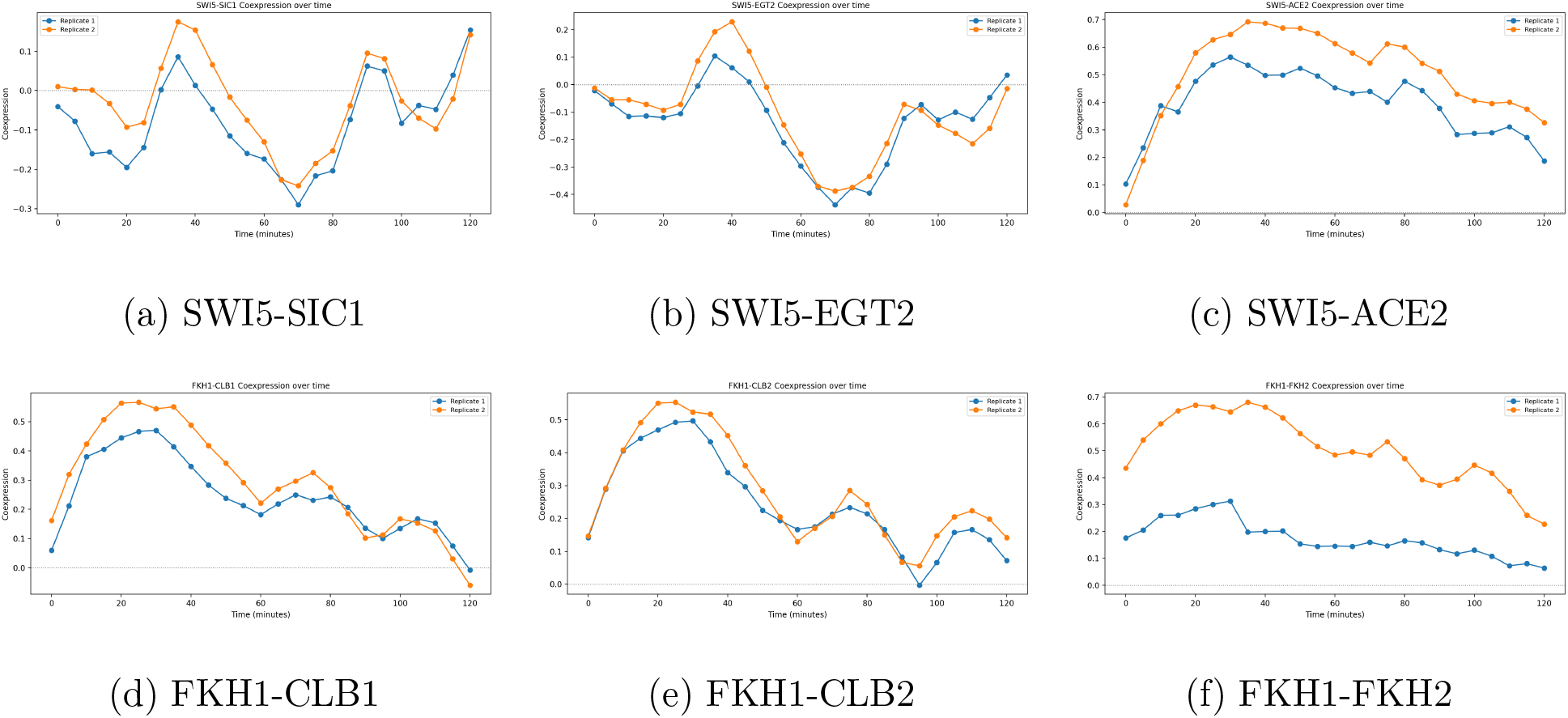
Time-varying PRISM coexpression edges for six representative gene pairs.

*HCM1* is a forkhead-family TF active during a specific window of the cell cycle, as characterized by the original study (Pramila et al. 2006). We test whether PRISM recovers the regulatory associations with *HCM1* that the original study identified. We used PRISM’s coexpression estimates to rank genes by their phase-contrast association with *HCM1* and performed gene set enrichment analysis against Gene Ontology Biological Process (Ashburner et al. 2000) terms independently in each replicate. We identified 10 pathways that were significant (at FDR 0.05) with concordant direction of enrichment in both replicates (Figure 5). Two of these, actin cytoskeleton organization and chitin biosynthetic process, correspond to the bud-morphogenesis functional category that Pramila et al. (2006) report among HCM1-regulated genes. The remaining pathways span translation (cytoplasmic translation, translational elongation), proteolysis, mating-pathway processes (cytogamy, karyogamy involved in conjugation), and a cluster of DNA-replication-related terms (DNA biosynthetic process, RNA-templated DNA biosynthetic process, DNA integration).

**Figure 5:**
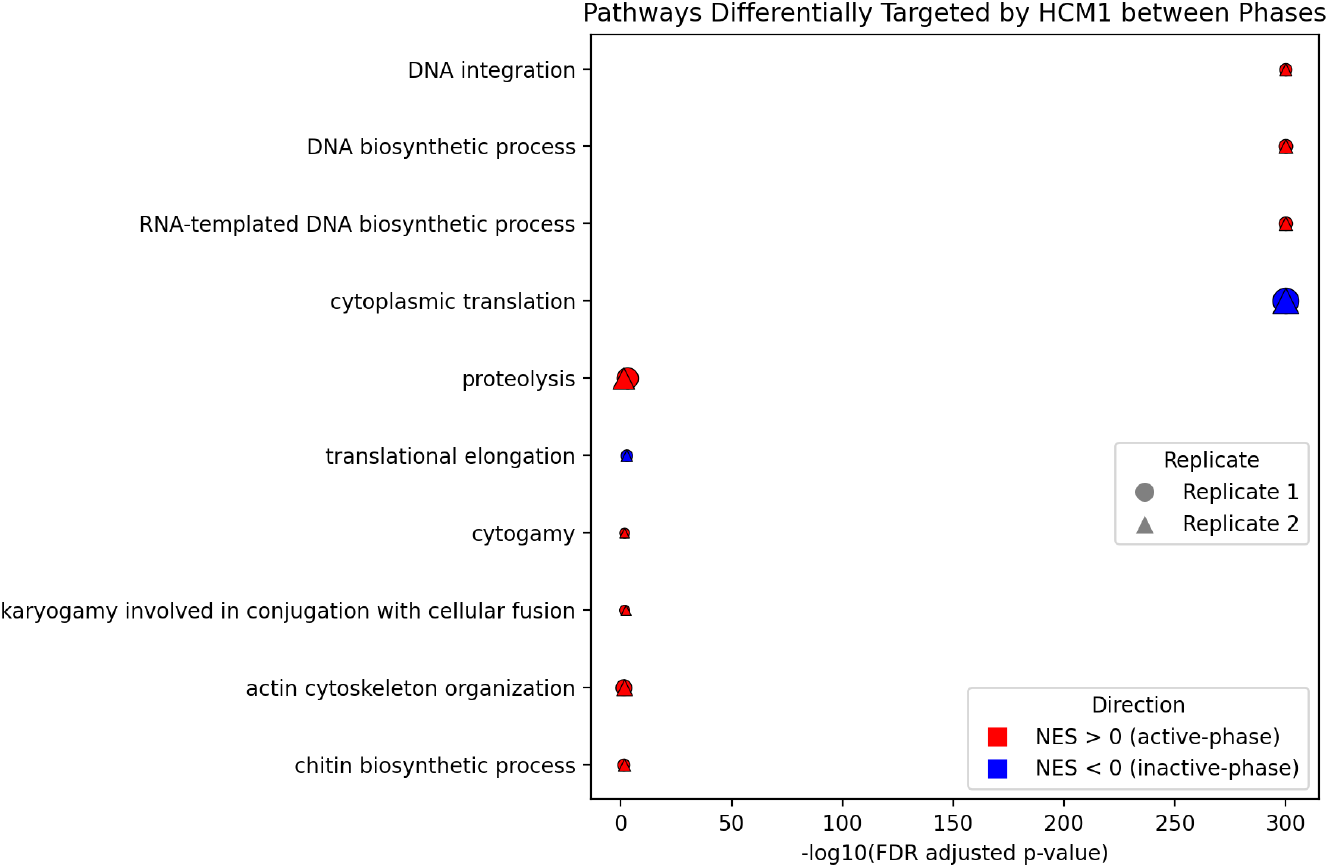
Pathways differentially targeted by HCM1 between active and inactive phases, validated across both biological replicates. Bubble size is proportional to gene set size, vertical axis shows pathway names and horizontal axis shows statistical significance in each replicate.

## 5 Application on Ovarian Cancer Data

Chromosomal instability caused by copy number alterations (CNA) play a major role in ovarian cancer (Network et al. 2011), where CNA amplification and deletion are known to induce altered expression across a majority of genes (Shao et al. 2019), with cis-effects (association between the CNA and expression levels of the same gene) dominating trans-effects (CNA-expression association across different genes) (Wang et al. 2011). TFs are regulatory proteins often driving such trans-effects, since a change in a TF’s own dosage can alter the expression of its downstream target genes (Consortium 2020). Ovarian tumors are molecularly heterogeneous, so the regulatory relationship between a given TF’s dosage and its targets’ expression may differ across patients. PRISM’s sample-specific networks are designed precisely for this setting, estimating each patient’s own coexpression structure rather than a single network pooled across the cohort. We study whether correlation between CNA in TF genes and the expression of their target genes, estimated by PRISM networks, is predictive of survival outcome in the ovarian cancer dataset from the Cancer Genome Atlas (TCGA-OV) (Weinstein et al. 2013).

We obtained RNA-seq gene expression, continuous copy number segment data, and phenotypic data via TCGAbiolinks (Colaprico et al. 2016). Copy number segments were mapped onto gene-level values for the TFs curated by (Lambert et al. 2018). Expression and copy number data were filtered, and matched at the patient level (full processing details are given in Supplementary Section S2), yielding a final combined matrix of 1,389 TFs (copy number) and 15,374 genes (expression) across 423 tumor samples, used as input to PRISM.

Copy number data are naturally non-Gaussian because a substantial fraction of samples are copy-neutral at any given gene, producing a point mass at the diploid value alongside a continuous tail for gains and losses. In TCGA-OV data, out of the 1,389 TFs, the Shapiro-Wilk test of normality was rejected for 1,386 (99.78%), at *α* = 0.05, and QQ-plots for the ten TFs with the highest CNA variance (Supplementary Figure S10) show the expected pattern: a flat plateau with a continuous tail. Therefore, we use the Gaussian copula variant of PRISM.

We ran PRISM with the “OAS” prior (Chen et al. 2010) on 423 samples. Consistent with cis-dominant regulation, cis network edges are overwhelmingly positive (98%), whereas trans edges are approximately symmetric about zero (Figure 6). To test whether trans-regulatory magnitude depends on TF motif presence in the target gene’s promoter (motif predictions from Marbach et al. (2016); available for 505 of 1,389 TFs), for each motif-covered TF, we tested whether its predicted targets were overrepresented among its largest-magnitude trans edges (one-sided Fisher’s exact test). Overrepresentation increased monotonically with a stricter magnitude threshold, reaching 113 of 505 TFs at the top 1% after FDR correction, while the analogous test in the opposite direction showed a diverging, decreasing trend, confirming a motif-specific effect rather than a generic association with edge magnitude (Supplementary Section S2). The five most significant TFs, RFX1, NR5A1, TP73, ALX3, and ESRRA, have all been documented in ovarian cancer (Issac et al. 2021, Lingasamy et al. 2026, Shoqafi et al. 2025, Wrzeszczynski et al. 2011, Salzman et al. 2011).

**Figure 6:**
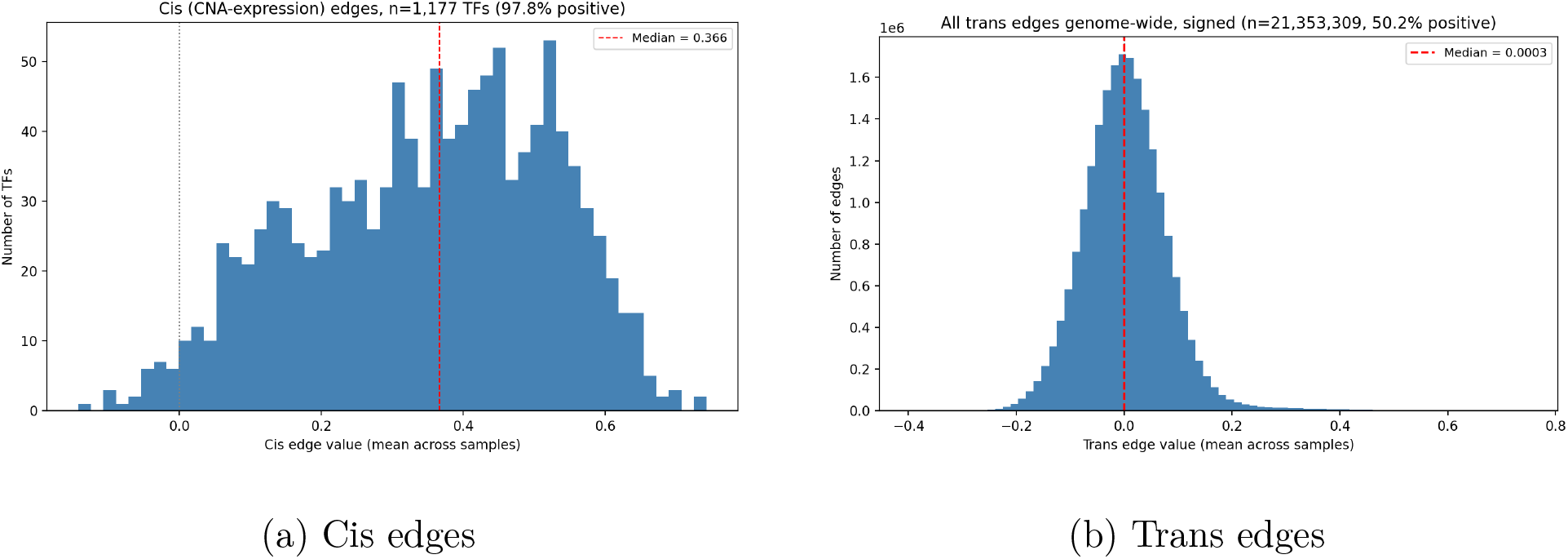
Distribution of cis and trans CNA-expression edges in TCGA-OV. Cis edges are overwhelmingly positive, whereas trans edges are approximately symmetric about zero.

Each of our CNA-expression networks derived by PRISM contains 21,354,486 total edges (1,177 cis and 21,353,309 trans). We performed edge-wise survival analysis using Cox proportional hazard models (Cox 1972) adjusting for patient age, race, and pathologic tumor stage, for all cis edges and the top 5% trans edges (1,067,665 edges) by variance across samples since a Cox model can only detect an association for edges that vary across patients. Genes whose CNA-driven dosage relationship with a target’s expression is predictive of survival are natural candidates for follow-up as potential therapeutic targets, since modulating such a gene’s expression, or the pathway it regulates, may influence patient outcome.

Among the top 50 trans edges (Figure 7, Kaplan-Meier curves for top ten in Supplementary Figure S11.) that were found to be significantly associated with survival, (FDR *<*0.05), the *HNF4G*-*GTF2H5* edge was found to have protective effects (hazard ratio *<* 1). HNF4G is a candidate transcriptional driver of chemoresistant tumor cells in high-grade serous ovarian cancer (Croft et al. 2024). *PAX8*, an established oncogenic TF in epithelial ovarian cancer (Di Palma & Zannini 2022), has a similar edge to *CDHR5* (hazard ratio *<*1). Among harmful edges (hazard ratio *>*1), *CACNB1* and *RSKR* were the most frequently recurring targets, each independently linked to ovarian cancer prognosis (Cai et al. 2021, Arechavaleta-Velasco et al. 2016). *ZIK1*, one of the TFs associated to *RSKR*, has separately been identified as differentially methylated in ovarian clear cell carcinoma (Li et al. 2024). However, all 20 TFs associated with *CACNB1* and 13 associated with *RSKR* lie within a single tandem zinc-finger cluster on chromosome 19 (spanning *∼*690 kb and 280 kb), with median pairwise CNA correlations of 0.97 and 0.99 and no documented binding motif for either target (Marbach et al. 2016), which is more consistent with one shared copy-number event than 33 independent regulatory associations. Individual literature support for a single TF in a cluster like this, such as *ZIK1*, does not establish it as the causal driver over its co-located, highly correlated neighbors without experimental validation (e.g., gene knockdown or CRISPR).

**Figure 7:**
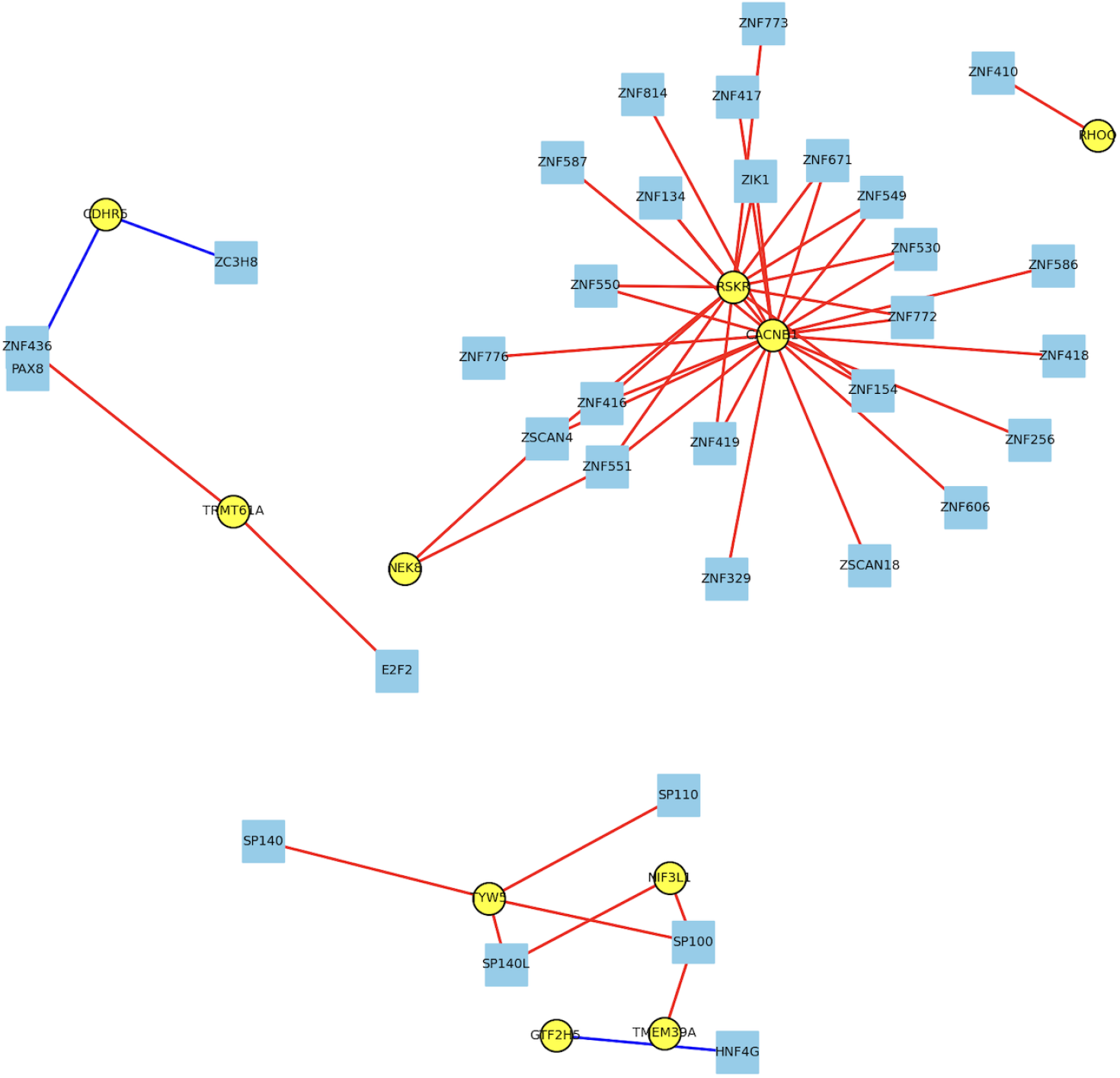
Network of the top 50 significant trans edges (FDR *<*0.05); squares=TF, circles=gene, red=harmful (HR> 1), blue=protective (HR <1).

## 6 Discussion

We introduced PRISM, an empirical Bayes framework for estimating sample-specific coexpression networks from cross-sectional and longitudinal multiomic data. PRISM imposes a conjugate prior on the sample-specific covariance matrices, whose parameters are estimated from the data thus borrowing strength from other samples and yielding a closed form posterior and edge-level uncertainty quantification. Relative to BONOBO (Saha et al. 2024), the closest existing framework, PRISM reduces computational cost from *O*(*n*^2^*g*^2^) to *O*(*ng*^2^) while matching BONOBO’s point-estimate accuracy throughout our simulations on Gaussian data. For non-Gaussian data, we observed a substantial advantage over BONOBO in uncertainty quantification. BONOBO which assumes an implicit Gaussian likelihood, showed severe undercoverage compared to PRISM which attained near-nominal coverage, even though the point estimates from both methods were comparable.

We applied PRISM to a longitudinal yeast cell-cycle dataset, where the estimated window-size hyperparameter and coexpression networks were reproducible across independent biological replicates, and PRISM recovered regulatory relationships and pathways independently documented in the cell-cycle literature. In a second real-data application, we used PRISM on paired copy-number and gene-expression data from ovarian cancer patients, recovering known biology such as cis-dominant regulatory patterns and a motif-dependent trans-regulatory signal. In addition, we identified novel interactions between TF copy number and target gene expression associated with survival outcome, including edges involving HNF4G and PAX8, TFs with independent evidence as, respectively, a candidate driver of chemoresistance and an established oncogenic regulator in ovarian cancer. These two applications, spanning distinct data modalities and study designs, demonstrate that PRISM recovers biologically meaningful, literature-consistent regulatory structure, and suggest that PRISM networks may recover gene regulatory mechanisms with potential clinical relevance.

PRISM has several limitations that we intend to address in future work. In presence of missing data, we assume a missing-completely-at-random mechanism, which is unlikely to hold exactly for single-cell data where lowly expressed genes are more prone to dropout. Extensions that incorporate an explicit missingness model (Sarkar & Stephens 2021) would be an interesting direction of future work, even though it might incur additional computational complexity due to the absence of a closed-form posterior update available here. In presence of structural population heterogeneity, PRISM requires either known group labels or a separate clustering step whose uncertainty is not propagated into the downstream network estimates. A hierarchical extension with mixture modeling might address this issue. For longitudinal data, PRISM selects a single window size shared across all genes, guaranteeing a positive semi-definite local prior by construction but forcing genes with different temporal autocorrelation to share one bandwidth. A natural next step would be to allow the window to vary smoothly over time while remaining shared across genes. Finally, the degrees-of-freedom calibration yields a common 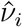 across individuals by default. In future work, we will explore possible extensions that enable individual-specific degrees of freedom parameters that capture how atypical a certain sample is compared to the rest of the population.

Because PRISM estimates a separate regulatory network for each patient rather than a single population-averaged network, it is naturally suited to settings where the same disease may be driven by different regulatory mechanisms in different individuals. As sample-specific multi-omic profiling becomes more widely available across cancer types and other heterogeneous diseases, we anticipate PRISM made available as open-source software, being used to identify which regulatory mechanisms are active in a given patient, a step toward informing individualized treatment strategies rather than population-level associations alone.

## 7 Data and Code Availability

The paper uses only publicly available datasets. Python code for PRISM implementation and additional code to reproduce the analysis presented in the paper are provided as supplementary files. Upon publication PRISM will be distributed as part of the SIGMAnet Python package: https://github.com/Enakshi-Saha/SIGMAnet.

## Supplementary Material for

Individual Sample-specific Gene Coexpression and Gene Regulatory Network Estimation with Empirical Bayes

### S1 Additional Proofs

#### Lemma 1.

*Suppose* 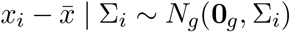 *and* Σ_*i*_ *∼ IW*(Ψ, *m*) *Then, marginalizing over* Σ_*i*_,

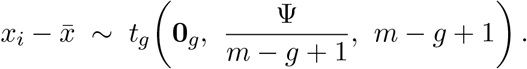

*In particular, under the prior* Σ_*i*_ *∼ IW*((*ν*_*i*_ *− g −* 1)*V, ν*_*i*_) *(i*.*e*. Ψ = (*ν*_*i*_ *− g −* 1)*V and m* = *ν*_*i*_),

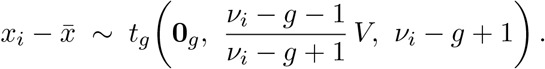

*Proof*. Let 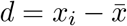. Multiplying the Gaussian likelihood and the inverse-Wishart density,

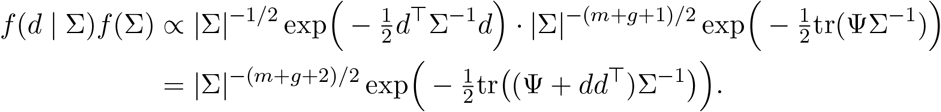

This is an *IW*(Ψ + *dd*^*⊤*^, *m* + 1) density up to a normalizing constant. Thus, integrating over Σ gives the reciprocal of that density’s normalizing constant:

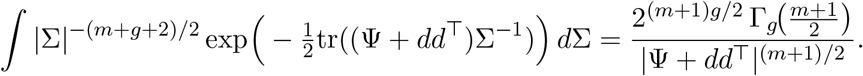

Combining with the normalizing constant of *f* (Σ) and the Gaussian constant (2*π*)^*−g/*2^,

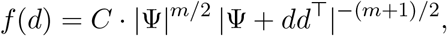

Using |Ψ + *dd*^*⊤*^| = |Ψ|(1 + *d*^*⊤*^Ψ^*−*1^*d*, we get

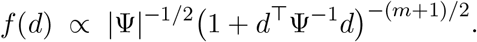

This is a *g*-dimensional multivariate-*t* density with location **0**_*g*_, sca le Ψ*/*(*m − g* + 1), and degrees of freedom *m − g* + 1. Hence *d ∼ t*_*g*_ (**0**_*g*_, Ψ*/*(*m − g* + 1), *m − g* + 1

#### Lemma 2.

*Under the model of Lemma 1, let Q*_*i*_(*V*) = *d*^*⊤*^*V* ^*−*1^*d and R*_*i*_ = *Q*_*i*_(*V*)*/g, where* 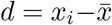.

*Then, for every ν*_*i*_, *R*_*i*_ *∼ c · F* (*g, ν*_*i*_ − *g* + 1), *where* 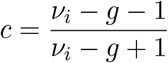 *and E*[*R*_*i*_] = 1.

*Proof*. By Lemma 1, *d ∼ t*_*g*_(**0**_*g*_, *cV, ν*^*\**^) with 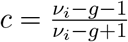 and *ν*^*\**^ = *ν*_*i*_ *− g* + 1.

For *d ∼ t*_*g*_(**0**_*g*_, Σ^*\**^, *ν*^*\**^), we have *d*^*⊤*^ (Σ^*\**^)^*−* 1^ *d/g ∼ F* (*g, ν*^*\**^). Thus, with Σ^*\**^ = *cV*,

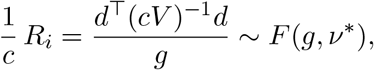

so *R*_*i*_ *∼ c · F* (*g, ν*^*\**^) = *c · F* (*g, ν*_*i*_ *− g* + 1).

Since 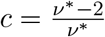 and 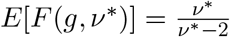 for *ν*^*\**^ *>* 2,

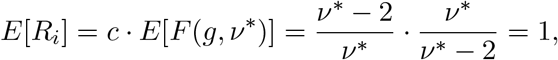

#### Lemma 3.

*Let* 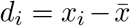 *denote the centered observation of individual i, let O* = *O*_*i*_ *and M* = *M*_*i*_ *denote its observed and missing coordinates, and let p* = |*O*| *a*(*nd q* = |*M* | = *g − p. Let d*_*O*_ *denote the sub-vector of d*_*i*_ *with coordinates in O. Suppose* Σ_*i*_ *∼ IW*_*g*_ (*ν − g −* 1)*V, ν*), *and that only d*_*O*_ *is observed, with*

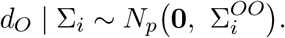

*Define*

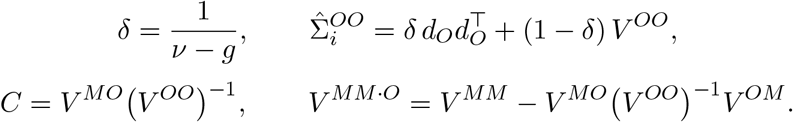

*Then the posterior means of the three blocks of* Σ_*i*_ *are*

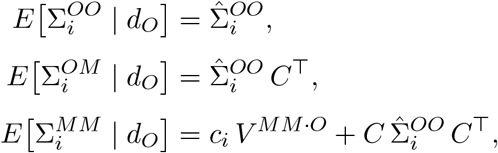

*where*

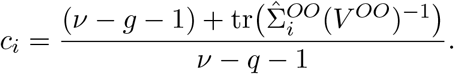

*In particular, with the imputed vector* 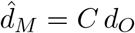,

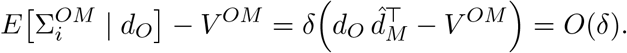

*Proof*. Throughout, we drop the subscript *i* and write Ψ = (*ν − g −* 1)*V* for the prior scale matrix.

Let *B* = (Σ^*OO*^) ^*−*1^Σ^*OM*^ and Σ^*MM·O*^ = Σ^*MM*^ *−* Σ^*MO*^ (Σ^*OO*^) ^*−*1^Σ^*OM*^. Under the inverse-Wishart prior, we get [Mardia et al., 2024].

a. Σ^*OO*^ *∼ IW*_*p*_ Ψ^*OO*^, *ν − q*.
b. Σ^*MM·O*^ *∼ IW*_*q*_ Ψ^*MM·O*^, *ν*, independently of Σ^*OO*^.
c. Given Σ^*OO*^ and Σ^*MM·O*^, *B* is matrix-normal with mean (Ψ^*OO*^)^*−*1^Ψ^*OM*^ = *C*^*⊤*^, row covariance (Ψ^*OO*^)^*−*1^ and column covariance Σ^*MM·O*^.

where the blocks and Schur complements of Ψ are defined analogously to those of *V*.

The likelihood of *d*_*O*_ depends on Σ only through Σ^*OO*^. Therefore,

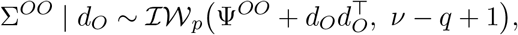

The posterior mean of Σ^*OO*^ is

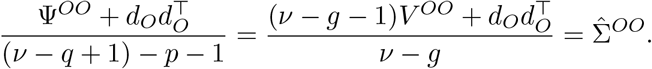

Since Σ^*OM*^ = Σ^*OO*^*B* and the two factors are independent,

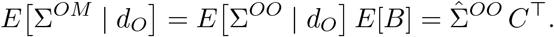

Expanding 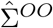 gives 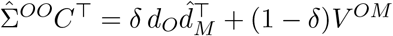, which proves the final claim.

Let Σ^*MM*^ = Σ^*MM·O*^ + *B*^*⊤*^Σ^*OO*^*B*. For a matrix-normal *A ∼ MN* (0, *U, W*) with row covariance *U* and column covariance *W, E*[*A*^*⊤*^*KA*] = tr(*KU*) *W*. Applying this to *B − C*^*⊤*^ gives

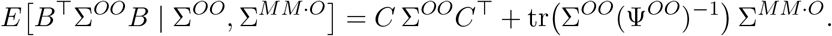

Taking expectations over the posterior gives

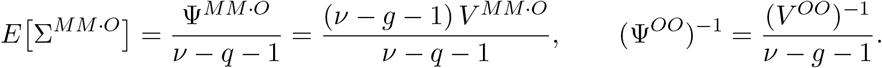

Thus,

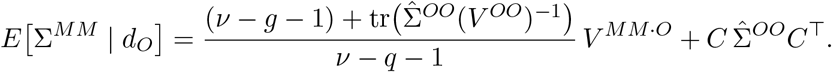

### S2 TCGA-OV Data Processing and Motif Overrepresentation Analysis

#### S2.1 TCGA-OV Data Processing

We obtained RNA-seq gene expression (GDC STAR-Counts pipeline), continuous copy number segment data (GDC Masked Copy Number Segment, i.e. segment-mean log2 ratios with germline variants removed), and phenotypic data, including survival, for TCGA-OV via TCGAbiolinks [Colaprico et al., 2016], restricting to primary tumor samples (426 expression samples, 601 copy number samples, 608 clinical records). Because copy number segments do not align with gene boundaries, we mapped segment-level values onto gene-level values for the 1,639 transcription factors curated by [Lambert et al., 2018], using genomic coordinates taken directly from the expression data’s own annotation to guarantee a consistent genome build (GRCh38) between the two modalities. For each sample, we overlapped that sample’s segments with each transcription factor’s coordinates. For genes spanning more than one segment, we computed a length-weighted average of the overlapping segments’ values, weighted by the width of overlap. Sixty-five transcription factors had no overlapping segment in any sample. Then we removed all TFs with at least one sample missing, yielding a final panel of 1,389 transcription factors with continuous copy number values.

We downloaded transcripts per million (TPM)-normalized gene expression data and retained genes with TPM *≥* 1 in at least 10% of samples and log-transformed the expression values (log(1 + TPM)). Ensembl gene identifiers were mapped to gene symbols using the expression data’s own annotation, the same annotation used to derive transcription factor coordinates for the copy number mapping above. Where multiple Ensembl identifiers mapped to the same symbol, we retained the identifier with the highest variance across samples, and further restricted to only protein-coding genes, yielding 15,374 genes. Expression and copy number samples were matched by truncating each dataset’s sample barcode to the patient level. Whereever this truncation produced duplicate columns within a modality (a small number of patients with multiple copy number aliquots), we retained the first occurrence and discarded the remainder. Restricting to samples present in both modalities yielded a final combined matrix of 16,763 omics variables (15,374 gene expression, 1,389 copy number) across 423 tumor samples, used as input to PRISM’s Gaussian copula variant.

For downstream survival analysis, 3 patients were excluded for missing pathologic tumor stage information and 1 additional patient for missing survival time, yielding a final set of 419 patients. Age was missing for 8 patients and imputed with the cohort median (59 years). Patients self-identified as White (n=363), Black (n=30), or another race (n=30). Because only one patient in this cohort was diagnosed at Stage I, we further combined Stage I and Stage II into a single category prior to model fitting, yielding a final stage distribution of Stage I-II (n=25), Stage III (n=329), and Stage IV (n=66).

#### S2.2 Motif Overrepresentation Analysis

For each of the 505 motif-covered TFs, we tested whether genes carrying a predicted motif for that TF were overrepresented among its largest-magnitude trans edges using a one-sided Fisher’s exact test. The remainder of the 1,389 TFs in our CNA panel lacked motif target predictions, reflecting the well-documented incompleteness of existing TF-target databases. Since a predicted motif indicates a likely direct binding site, CNA-driven dosage effects on a transcription factor should propagate more strongly to targets with such a site than to targets without one.

The number of TFs reaching significance after multiple-testing correction (BH-FDR *<*0.05) increased monotonically as the magnitude threshold was tightened: 55 of 505 TFs (10.9%) among the top 10% of edges by magnitude, 61 (12.1%) among the top 5%, and 113 (22.4%) among the top 1%, indicating that motif-dependent enrichment concentrates specifically among the most extreme trans effects. Testing the opposite direction (motif-absent genes overrepresented among top edges) at the same thresholds yielded a pattern that diverged: 29 TFs (5.7%) at the top 10%, 20 (4.0%) at the top 5%, and only 18 (3.6%) at the top 1%, decreasing as the motif-present direction increased. This divergence is consistent with the motif-present enrichment reflecting a genuine, motif-specific effect rather than a generic tendency for extreme trans edges to co-occur with any fixed gene subset.

### S3 Supplementary Figures and Tables

**Figure S1:**
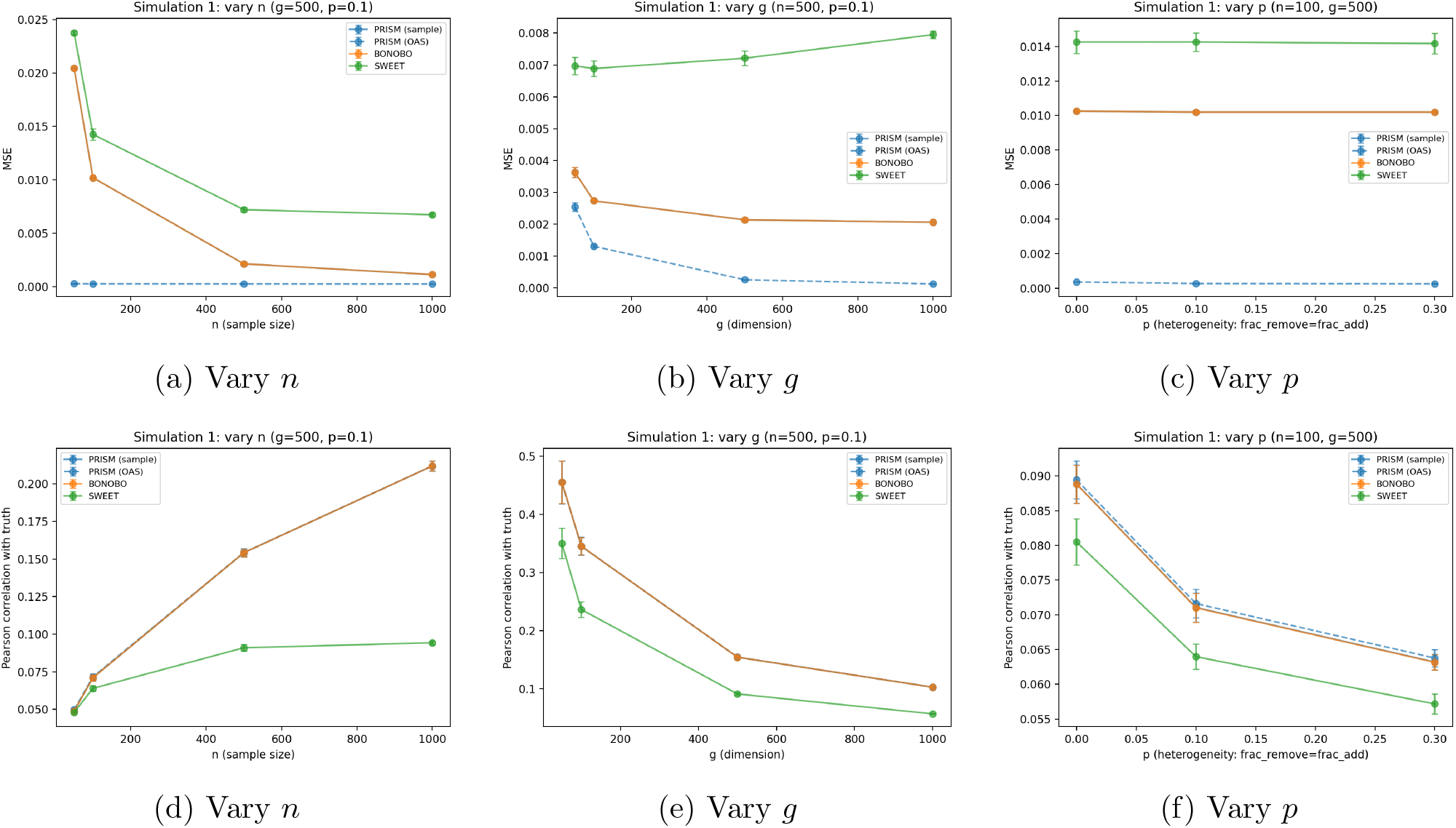
Simulation 1, supplementary metrics: MSE (top row) and Pearson correlation with the true correlation matrix (bottom row). PRISM’s Pearson correlation with the truth closely coincides with BONOBO’s (panels d–f), indicating comparable recovery of the true correlation structure. However, PRISM’s MSE is near zero throughout, substantially lower than BONOBO’s and SWEET’s. Since Pearson correlation does not show a corresponding PRISM advantage, this MSE gap reflects PRISM’s shrinkage producing estimates of smaller overall magnitude rather than a proportionally better edge recovery. Pearson correlation is low across all methods, reflecting the large proportion of true non-edges in the sparse ground truth.

**Figure S2:**
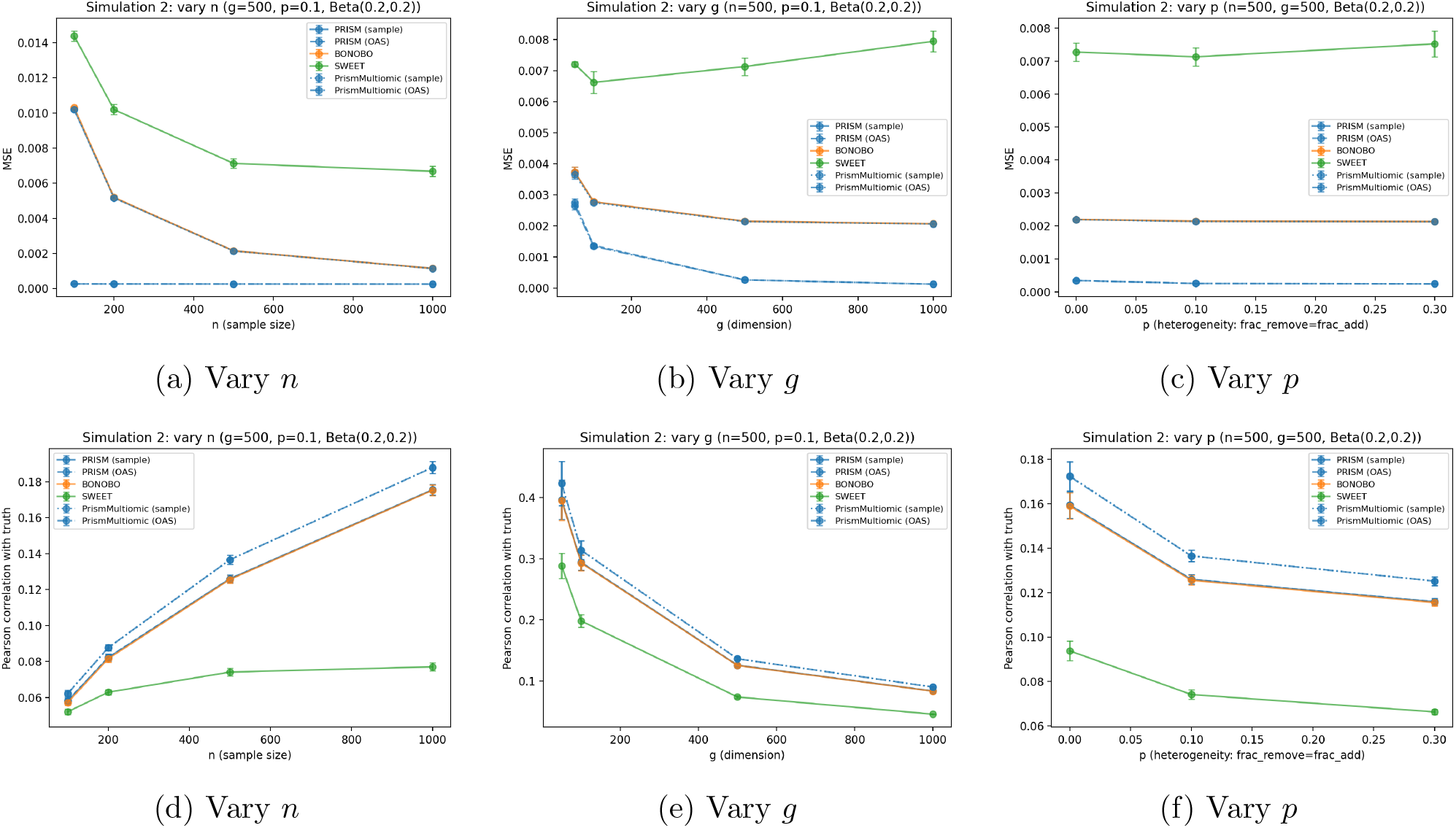
Simulation 2, supplementary metrics: MSE (top row) and Pearson correlation with the true correlation matrix (bottom row). As in Simulation 1, PRISM’s MSE (both prior variants, and their nonparanormal counterparts) is near zero throughout, substantially lower than BONOBO’s and SWEET’s. Because this is not matched by a correspondingly larger gap in Pearson correlation, it again reflects smaller estimate magnitude from shrinkage rather than proportionally better edge recovery. In Pearson correlation, the nonparanormal PRISM variants modestly but consistently exceed standard PRISM, most visibly at larger *n* (panel d). As in Simulation 1, Pearson correlation is low across all methods, reflecting the large proportion of true non-edges in ground truth.

**Figure S3:**
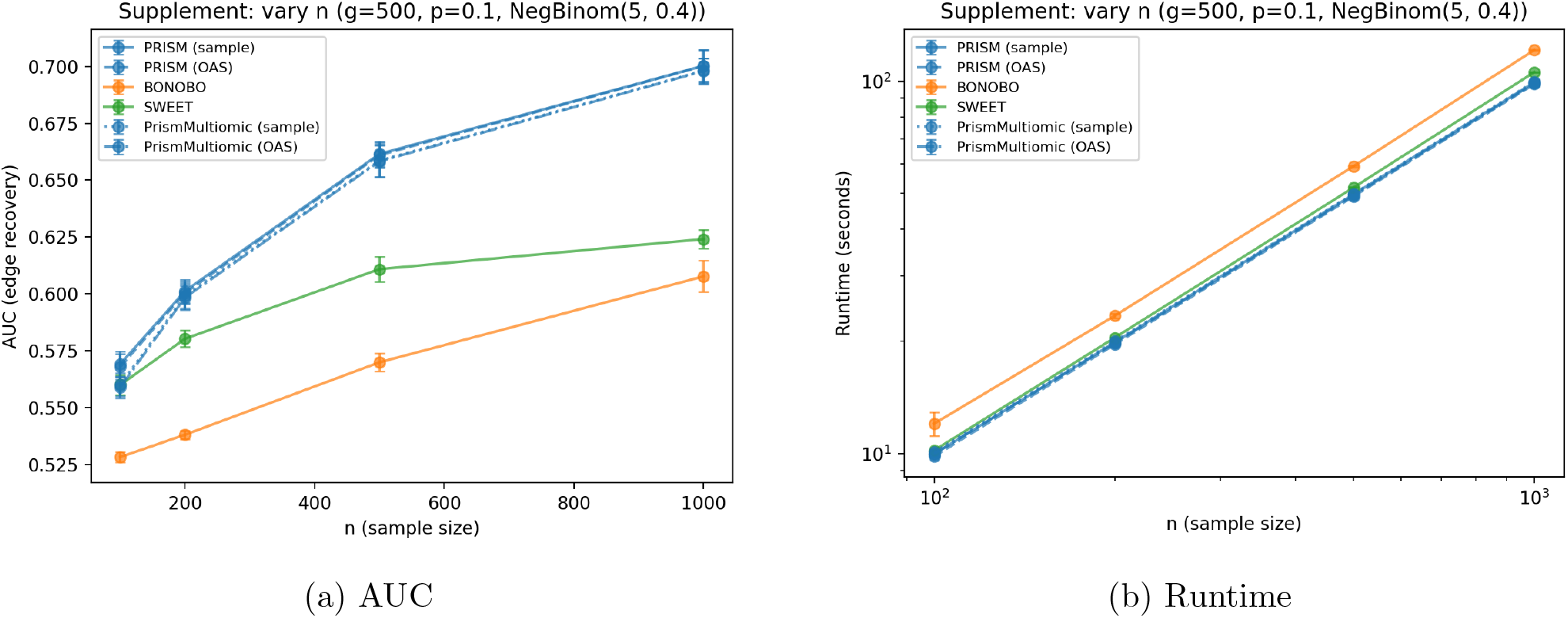
Supplementary check with Negative Binomial (CNV-like, tie-heavy count) marginals: edge-recovery AUC and runtime as a function of sample size *n*, fixed *g* = 500, *p* = 0.1. All PRISM variants (standard and nonparanormal, both prior choices) closely coincide in AUC across the full range of *n*, indicating no detectable benefit of the nonparanormal transform under this tie-heavy marginal distribution, unlike the Beta-marginal case in the main text; this is consistent with rank-based Gaussianization being less effective when many observations share identical values. Unlike the Beta and Gaussian settings, BONOBO underperforms SWEET in AUC here across all sample sizes. Runtime patterns are consistent with the other simulations: BONOBO is slowest and scales most steeply with *n*, while the PRISM variants remain close to SWEET.

**Figure S4:**
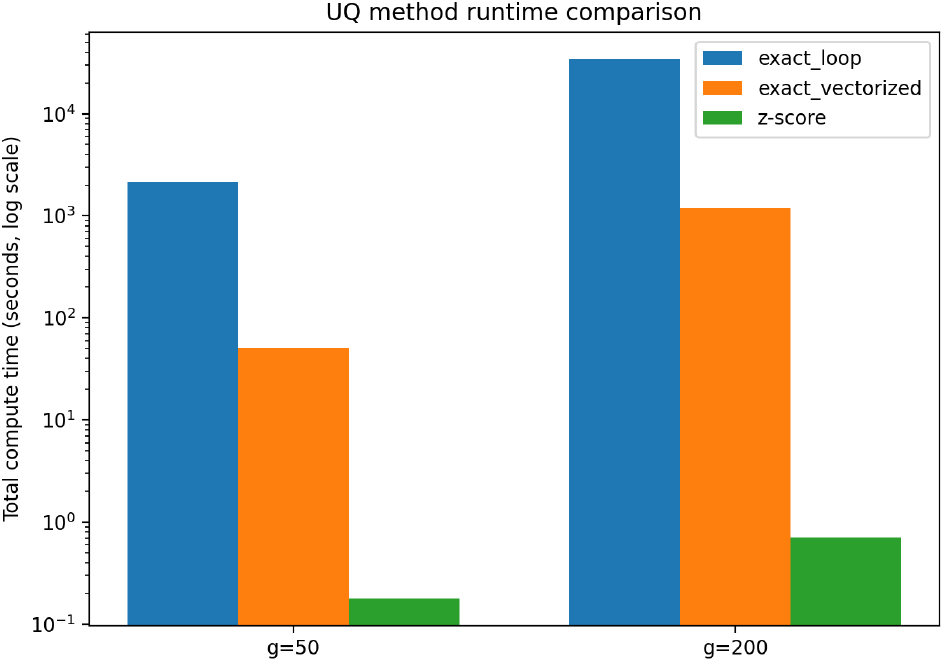
Runtime comparison of PRISM’s three edge-level uncertainty quantification approaches (exact, looped; exact, vectorized; z-score) at *n* = 100, *g* ∈ {50, 200}, extrapolated to 5 replicates *×* 100 samples. The looped method’s runtime is extrapolated from a subsampled set of edges rather than measured on the full edge set; the vectorized and z-score methods are measured directly on all edges.

**Figure S5:**
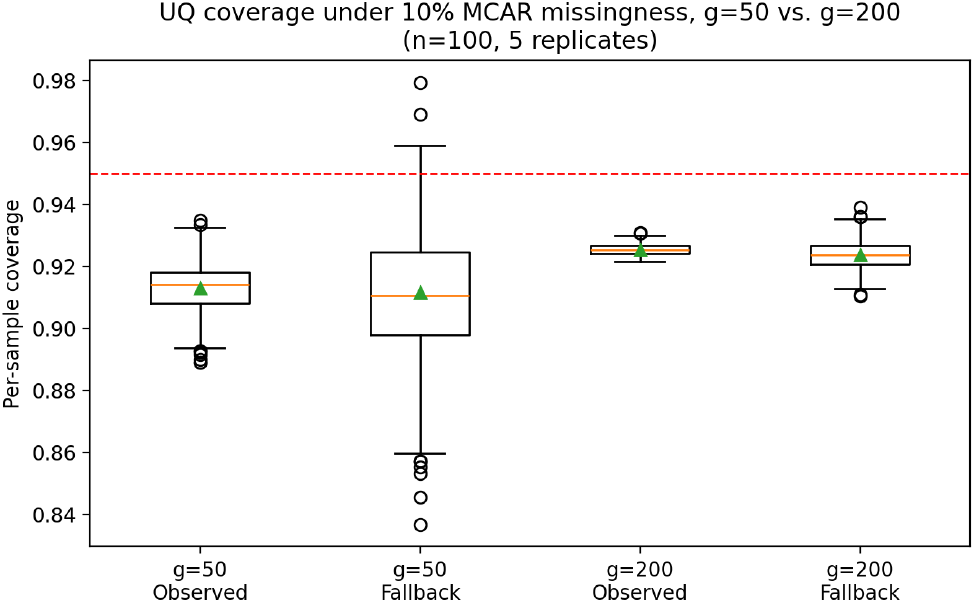
Per-sample coverage under 10% MCAR missingness, split by observed vs. fallback edges, at *g* = 50 and *g* = 200 (*n* = 100, 5 replicates). Dashed line: nominal 95% target.

**Figure S6:**
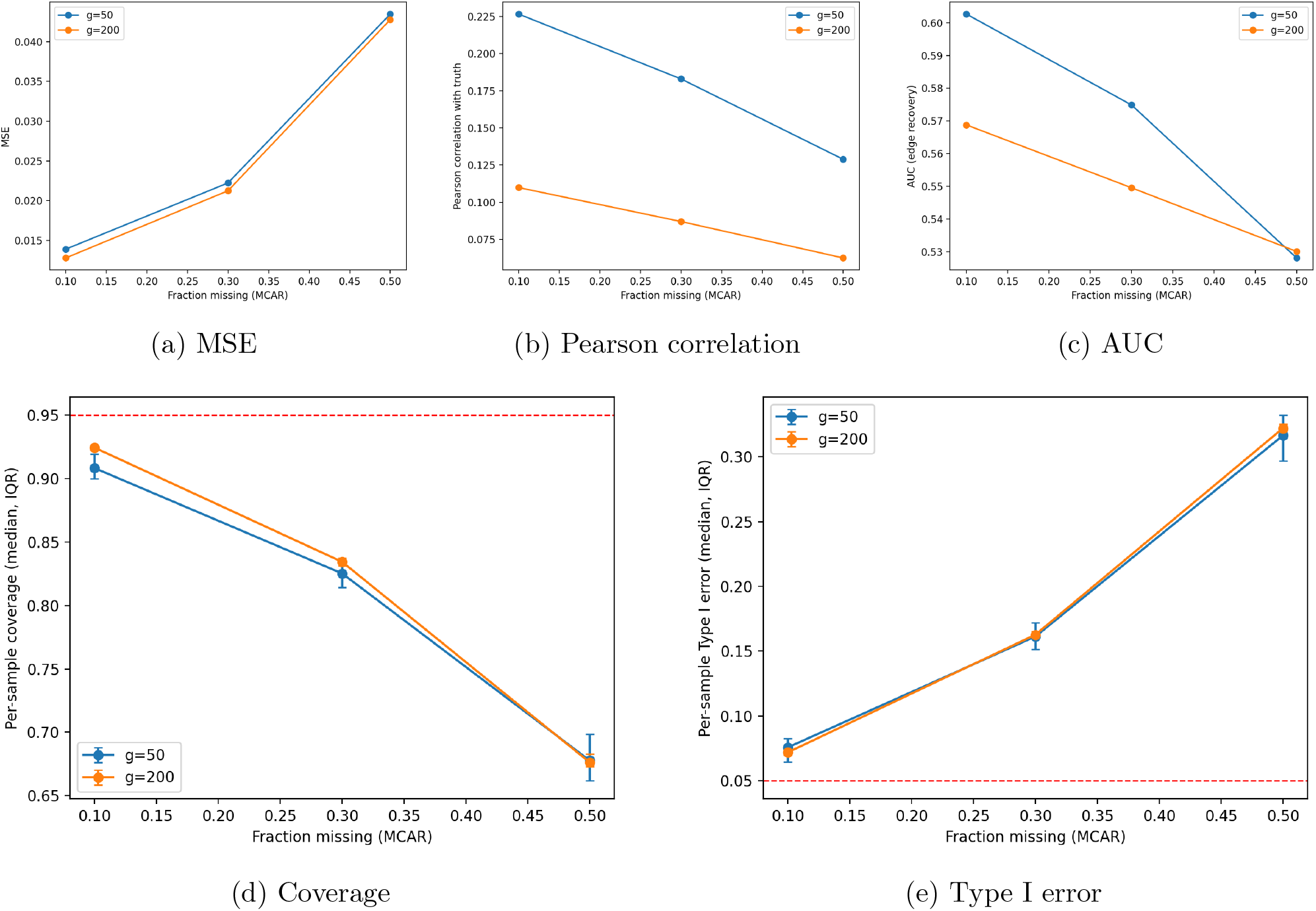
Point-estimate accuracy (top row) and edge-level UQ calibration (bottom row) as a function of MCAR missingness fraction (10–50%), for *g* = 50 and *g* = 200 (*n* = 100, 5 replicates). Dashed lines in (d)/(e): nominal targets (coverage = 0.95, Type I error = 0.05).

**Figure S7:**
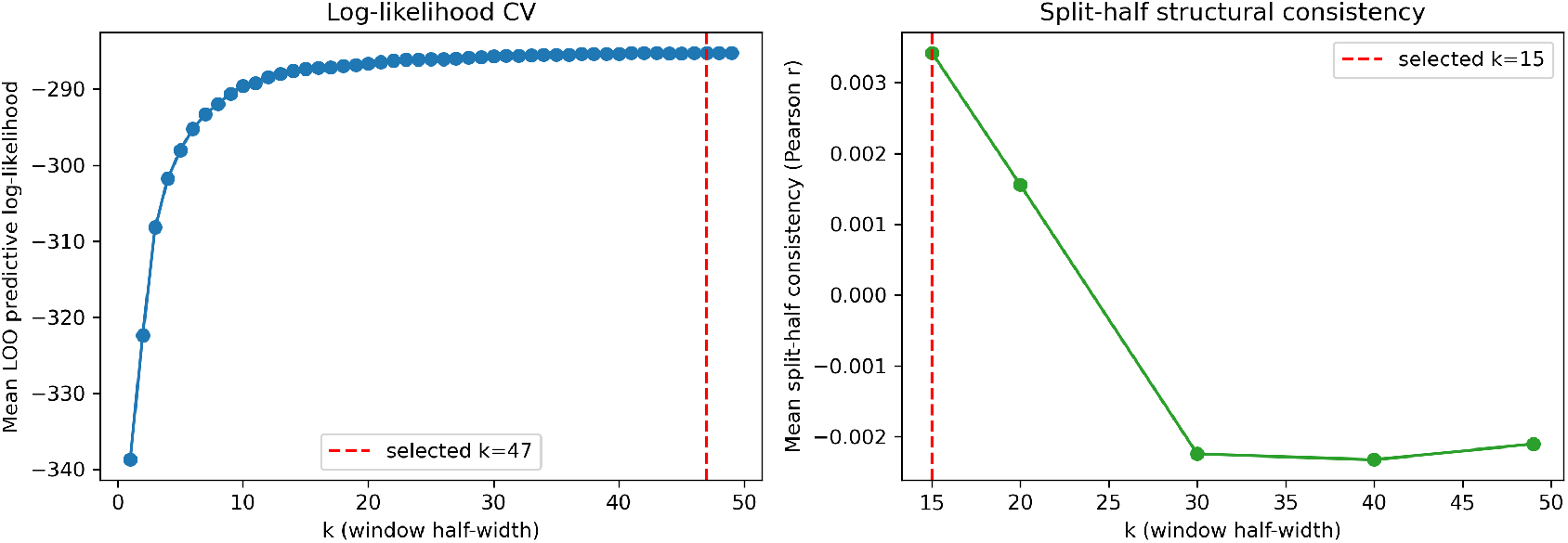
Log-likelihood cross-validation (left) vs. split-half structural consistency (right), stepwise drift, *g* = 200, *n* = 100. Log-likelihood CV selects *k* = 47; split-half selects *k* = 15, closer to the oracle recovery-optimal window (Table S2).

**Figure S8:**
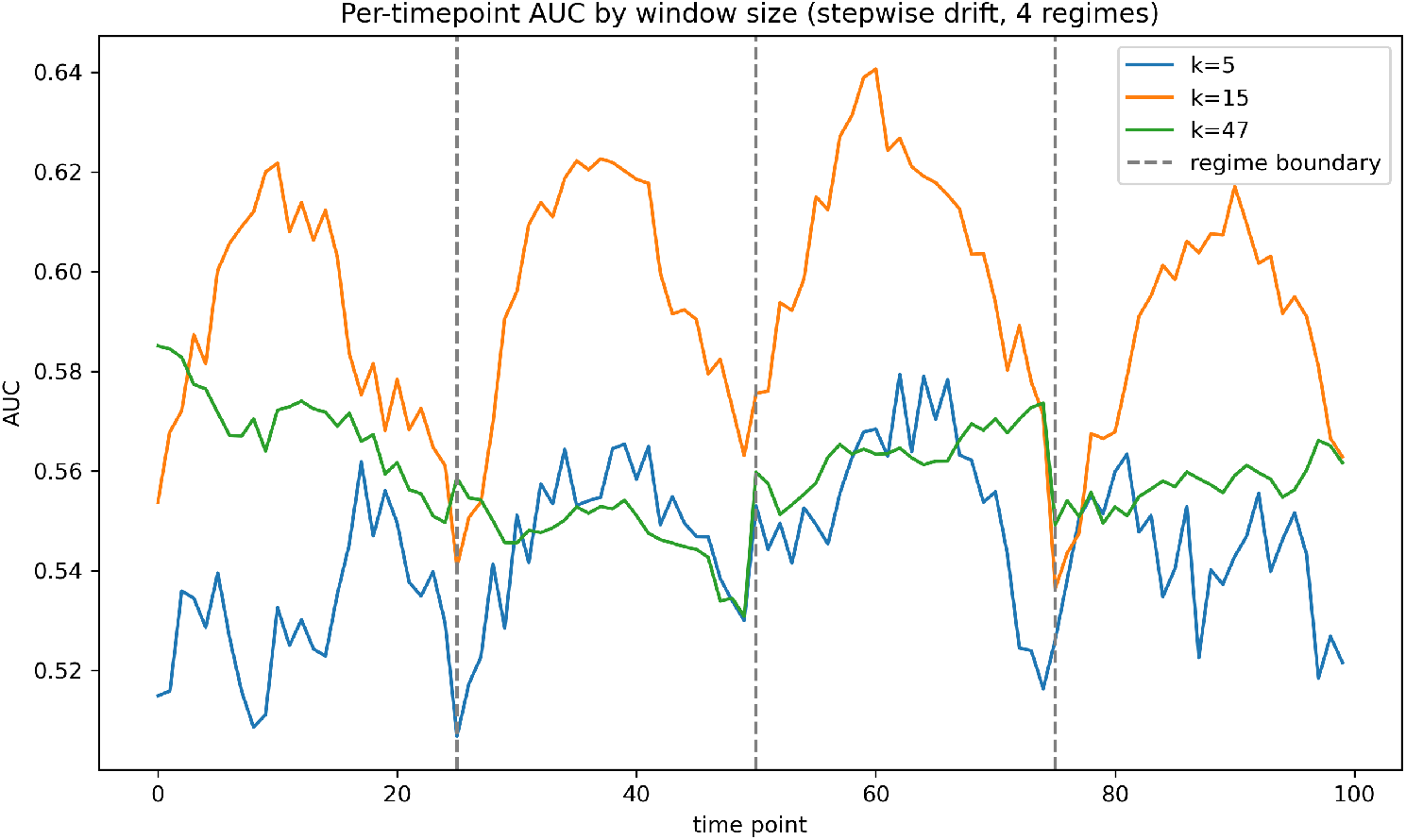
Per-timepoint AUC under stepwise drift for three representative window half-widths (*k* = 5, 20, 49), *g* = 200, *n* = 100. Dashed lines mark the three regime boundaries (*t ≈* 25, 50, 75). The moderate window (*k* = 20) achieves the highest AUC for time points well within a single regime but degrades sharply at each boundary; the largest window (*k* = 49) is more uniform but never reaches *k* = 20’s peak accuracy; the smallest window (*k* = 5) underperforms throughout.

**Figure S9:**
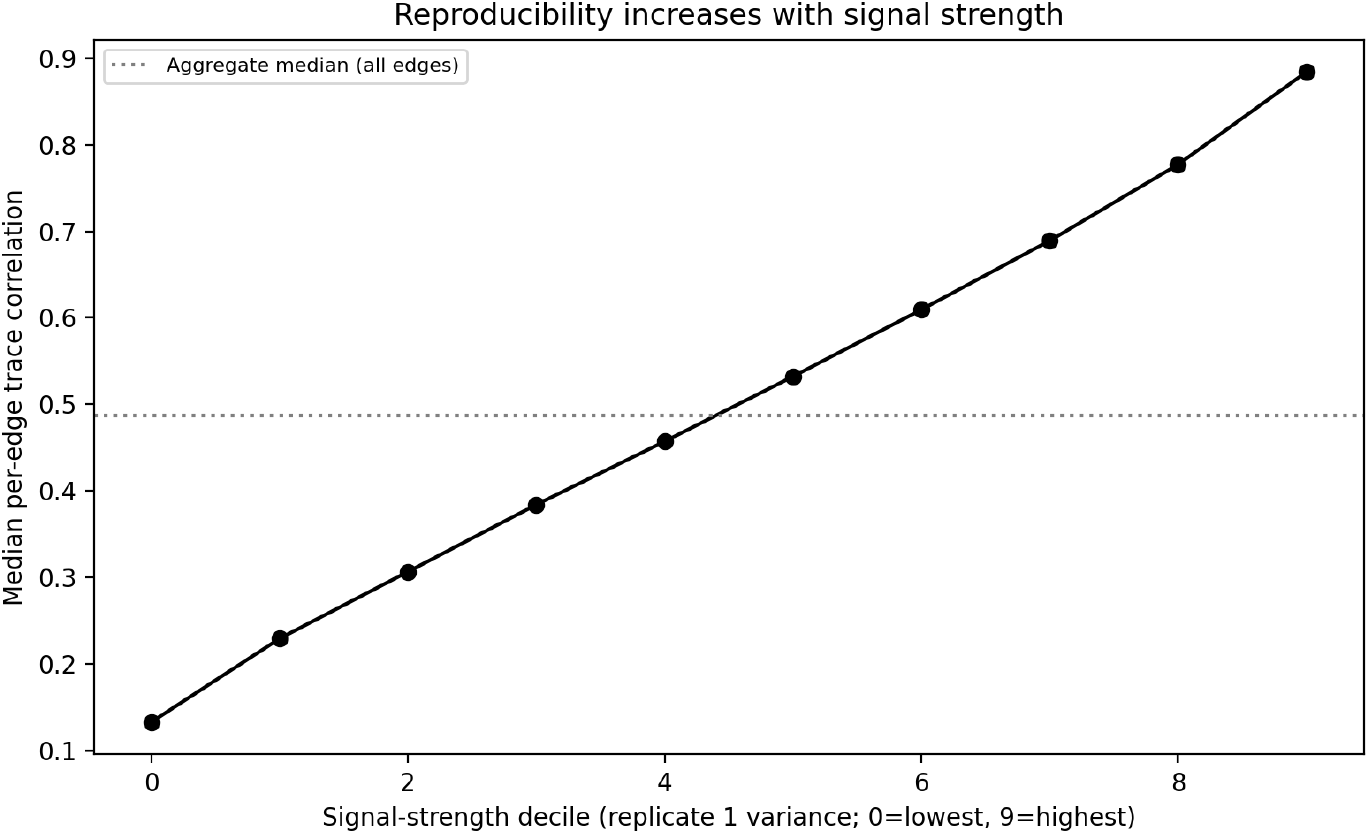
Reproducibility of PRISM coexpression edges between biological replicates in yeast cell cycle data. Median per-edge correlation increases monotonically with signal strength (variance decile), from 0.13 in the lowest decile to 0.88 in the highest decile.

**Figure S10:**
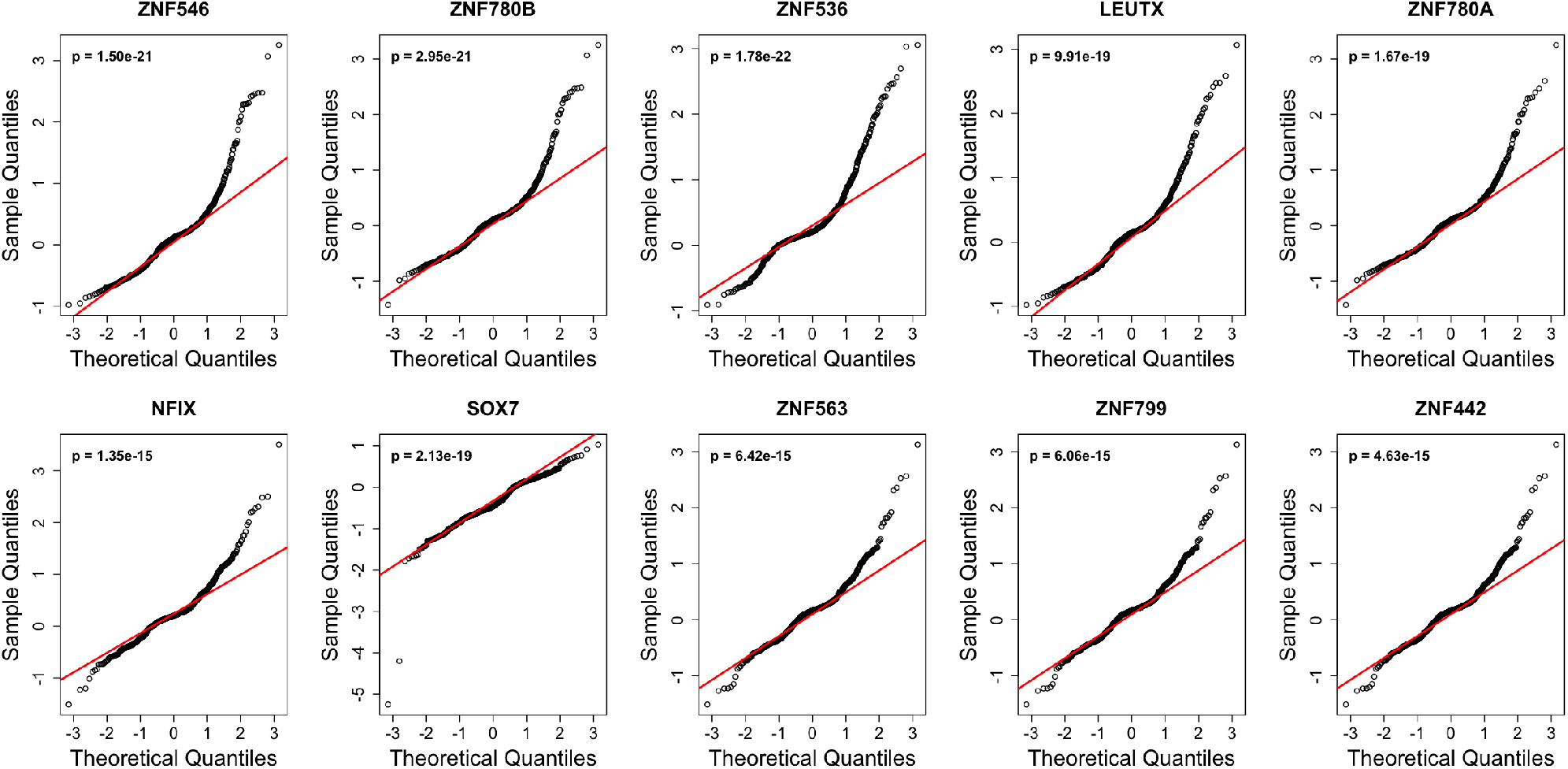
QQ-plots against a standard normal distribution for the ten transcription factors with the highest CNA variance in TCGA Ovarian Cancer data, with the Shapiro–Wilk test *p*-value reported in each panel. All ten show a clear departure from normality.

**Figure S11:**
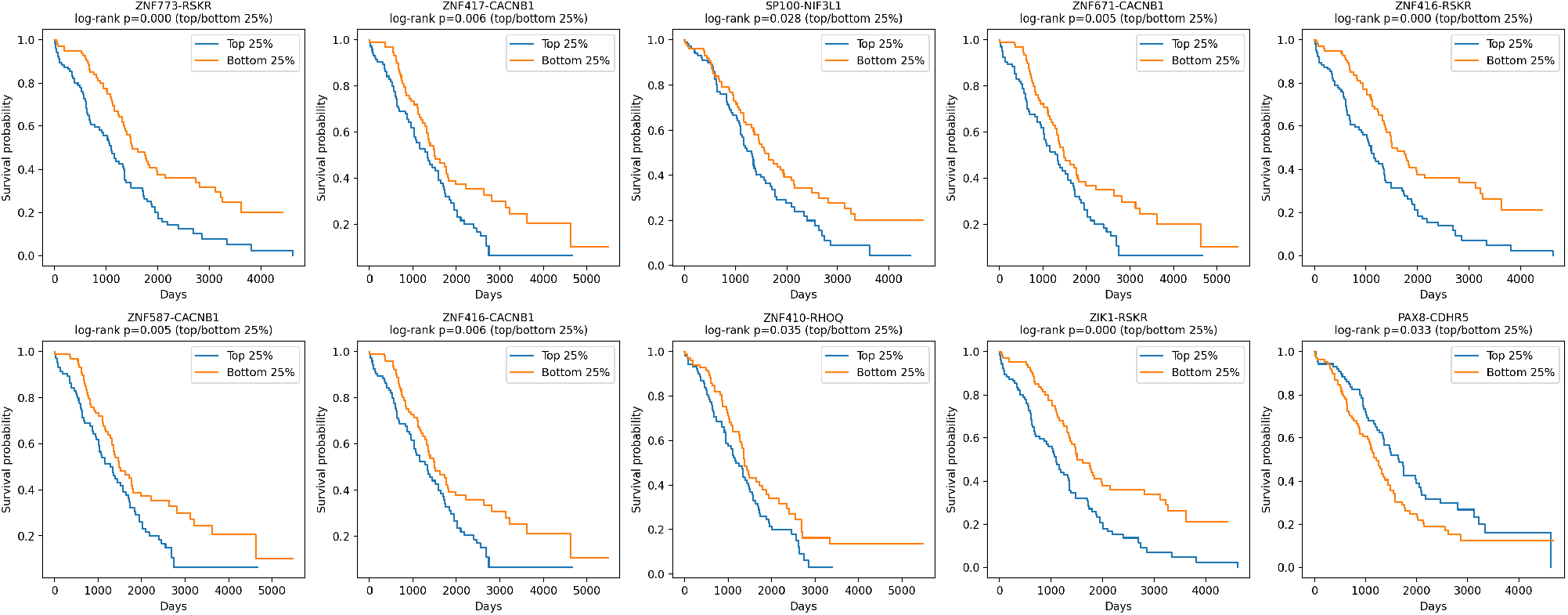
Kaplan-Meier curves for ten representative significant trans edges, dichotomized by top 25% and bottom 25% of edge value (unadjusted log-rank test). The Cox models reported in main text adjust for age, race, and stage and use the full continuous edge value.

**Table S1:** Per-sample coverage and Type I error (median [IQR], pooled across all evaluated individuals and 5 simulation replicates, *n* = 100) comparing PRISM’s three edge-level uncertainty quantification approaches: exact posterior sampling (looped and vectorized implementations, mathematically identical) and the closed-form z-score approximation, across dimension *g* and population structure. Nominal targets: coverage = 0.95, Type I error = 0.05.

| $g$ | Population | Method | Coverage | Type I Error |
| --- | --- | --- | --- | --- |
| 50 | Homogeneous | Exact, looped | 0.940 [0.930, 0.955] | 0.055 [0.043, 0.067] |
|  |  | Exact, vectorized | 0.944 [0.936, 0.947] | 0.054 [0.049, 0.062] |
|  |  | Z-score | 0.961 [0.955, 0.961] | 0.041 [0.039, 0.046] |
| 50 | Mixture (2 groups) | Exact, looped | 0.925 [0.910, 0.935] | 0.054 [0.042, 0.066] |
|  |  | Exact, vectorized | 0.925 [0.922, 0.927] | 0.053 [0.050, 0.057] |
|  |  | Z-score | 0.937 [0.936, 0.941] | 0.045 [0.043, 0.048] |
| 200 | Homogeneous | Exact, looped | 0.945 [0.930, 0.955] | 0.056 [0.041, 0.067] |
|  |  | Exact, vectorized | 0.942 [0.941, 0.943] | 0.056 [0.055, 0.057] |
|  |  | Z-score | 0.956 [0.956, 0.957] | 0.044 [0.043, 0.045] |
| 200 | Mixture (2 groups) | Exact, looped | 0.935 [0.925, 0.950] | 0.056 [0.046, 0.069] |
|  |  | Exact, vectorized | 0.936 [0.935, 0.937] | 0.058 [0.057, 0.059] |
|  |  | Z-score | 0.951 [0.950, 0.951] | 0.046 [0.046, 0.046] |

**Table S2:**
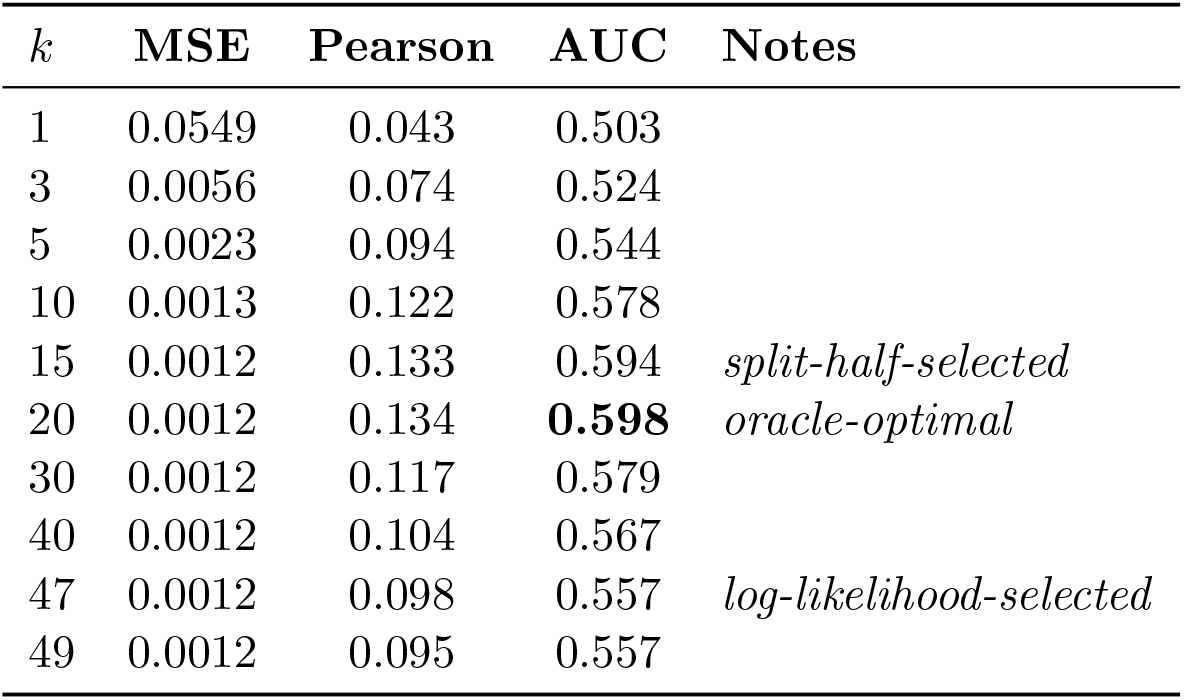
Edge-recovery accuracy as a function of window half-width *k*, for the stepwise drift (*g* = 200, *n* = 100, single dataset) example. AUC and Pearson correlation peak at *k* = 20; MSE plateaus for *k ≥* 15. The log-likelihood-selected window (*k* = 47) and split-half-selected window (*k* = 15) are marked. MSE is reported to higher precision than the other tables in this paper, since the compared methods differ only at this finer scale.

| $k$ | MSE | Pearson | AUC | Notes |
| --- | --- | --- | --- | --- |
| 1 | 0.0549 | 0.043 | 0.503 |  |
| 3 | 0.0056 | 0.074 | 0.524 |  |
| 5 | 0.0023 | 0.094 | 0.544 |  |
| 10 | 0.0013 | 0.122 | 0.578 |  |
| 15 | 0.0012 | 0.133 | 0.594 | *split-half-selected* |
| 20 | 0.0012 | 0.134 | **0.598** | *oracle-optimal* |
| 30 | 0.0012 | 0.117 | 0.579 |  |
| 40 | 0.0012 | 0.104 | 0.567 |  |
| 47 | 0.0012 | 0.098 | 0.557 | *log-likelihood-selected* |
| 49 | 0.0012 | 0.095 | 0.557 |  |

**Table S3:** Supplementary results for longitudianl data (simulation 4): MSE and Pearson correlation, mean *±* SD across 5 replicates, *g* = 200, *n* = 100. MSE is reported to higher precision than the other tables in this paper, since the compared methods differ only at this finer scale.

| Drift | Method | MSE | Pearson |
| --- | --- | --- | --- |
| Linear | PrismLongitudinal | $0.000759 \pm 0.000024$ | $0.205 \pm 0.005$ |
| | Naive PRISM | $0.000758 \pm 0.000023$ | $0.206 \pm 0.005$ |
| | Rolling window | $0.014110 \pm 0.000110$ | $0.204 \pm 0.005$ |
| Stepwise | PrismLongitudinal | $0.001218 \pm 0.000018$ | $0.134 \pm 0.004$ |
| | Naive PRISM | $0.001182 \pm 0.000021$ | $0.084 \pm 0.003$ |
| | Rolling window | $0.037460 \pm 0.000060$ | $0.132 \pm 0.008$ |

### S4 Downstream Application of PRISM: Individual-Specific Gene Regulatory Network Inference

The coexpression matrices estimated by PRISM for each individual are compatible with existing methods for gene regulatory network (GRN) inference that take a coexpression prior as input, such as PANDA [Glass et al., 2013] and OTTER [Weighill et al., 2021], following the same usage pattern established for BONOBO [Saha et al., 2024]. BONOBO’s own evaluation [Saha et al., 2024] demonstrates that coexpression matrices of this form, used as PANDA input, recover biologically meaningful, sample-specific regulatory structure. Since PRISM’s output has the same form (a symmetric, sample-specific gene-by-gene correlation matrix) and comparable accuracy to BONOBO’s, we do not repeat this evaluation here. We provide this functionality in the PRISM software.

